# Dinoflagellates with relic endosymbiont nuclei as novel models for elucidating organellogenesis

**DOI:** 10.1101/702274

**Authors:** Chihiro Sarai, Goro Tanifuji, Takuro Nakayama, Ryoma Kamikawa, Kazuya Takahashi, Hideaki Miyashita, Ken-ichiro Ishida, Mitsunori Iwataki, Yuji Inagaki

## Abstract

Nucleomorphs are relic endosymbiont nuclei so far found only in two algal groups, cryptophytes and chlorarachniophytes, which have been studied to model the evolutionary process integrating an endosymbiont alga into be a host-governed plastid (organellogenesis). Nevertheless, past studies suggested that DNA transfer from the endosymbiont to host nuclei had already ceased in both cryptophytes and chlorarachniophytes, implying that the organellogenesis at the genetic level has been completed in the two systems. Moreover, we have yet to pinpoint the closest free-living relative of the endosymbiotic alga engulfed by the ancestral chlorarachniophyte or cryptophyte, making difficult to infer how organellogenesis altered the endosymbiont genome. To counter the above issues, we need novel nucleomorph-bearing algae, in which from-endosymbiont-to-host DNA transfer is on-going and of which endosymbiont/plastid origins can be inferred at a fine taxonomic scale. Here, we report two previously undescribed dinoflagellates, strains MGD and TGD, with green algal endosymbionts enclosing plastids as well as relic nuclei (nucleomorphs). We provide the evidence for the presence of DNA in the two nucleomorphs and transfer of endosymbiont genes to the host (dinoflagellate) genomes. Furthermore, DNA transfer between the host and endosymbiont nuclei was found to be in progress in both MGD and TGD systems. Phylogenetic analyses successfully resolved the origins of the endosymbionts at the genus level. Combined, we conclude that the host-endosymbiont integration in MGD/TGD is less advanced than that in cryptophytes/chrorarachniophytes, and propose the two dinoflagellates as new models for elucidating organellogenesis.

## Introduction

The transformation of a free-living photosynthetic organism to the plastid through endosymbiosis occurred multiple times in eukaryotic evolution. The first plastid was most likely established through ‘primary endosymbiosis’ between a cyanobacterium and the common ancestor of red, glaucophyte, and green algae (plus descendants of green algae, i.e. land plants) (1). The plastids in the three lineages described above are the direct descendants of the cyanobacterial endosymbiont, and designated as ‘primary plastids.’ After major eukaryotic lineages were diverged, some heterotrophs turned into phototrophs by acquiring ‘secondary plastids’ through algal endosymbionts bearing primary plastids (secondary endosymbioses). Secondary endosymbioses most likely occurred multiple times in eukaryotic evolution, as the host lineages bearing secondary plastids (so-called complex algae) are distantly related to one another (2, 3). In addition, the origins of secondary plastids vary among complex algae—some possess red alga-derived plastids, while others possess green alga-derived plastids, strongly arguing that the two types of secondary plastids were established through separate (at least two) endosymbiotic events (2, 3).

The evolutionary process integrating an endosymbiont into the host cell (organellogenesis) has yet to be fully understood. Nevertheless, genomic data from diverse eukaryotic lineages indicated that endosymbiont genomes should have lost a massive number of genes that were dispensable for intracellular/endosymbiotic lifestyles (4). It is most likely that the reduction of endosymbiont genomes and integration of the endosymbiont into the host progressed simultaneously during organellogenesis (4). The reductive process worked on endosymbiont genomes seemingly has a tight correlation with genome G + C content, as reduced endosymbiont genomes are commonly poor in G and C (5–7). To interlock the host and endosymbiont metabolically and genetically, we regard transfer of endosymbiont genes to the host nuclear genome (endosymbiotic gene transfer or EGT), coupled with the inventions of host’s machineries that enable to express the transferred genes and target the gene products to the original compartment, as critical (8). Nevertheless, the precise process that enables organellogenesis still remains unclear.

In most of complex algae, no intracellular structure of the endosymbiotic algae was left except plastids. However, only cryptophytes and chlorarachniophytes are known to retain nucleomorphs, the relic nuclei of their algal endosymbionts (9, 10). As the two complex algae bearing nucleomorphs possess the morphological characteristics that have been lost from others, they were anticipated to provide clues to understand the detailed process of organellogenesis (7). In this regard, the genomic data of both nuclei and nucleomorphs, as well as transcriptomic and proteomic data have been accumulated for cryptophytes and chlorarachniophytes (11–13), and genetic transformation was established for a chlorarachniophyte species (14). It has been defined a red alga and a ulvophyte green alga as the origins of cryptophyte and chlorarachniophyte plastids, respectively (15–17), but an even recent multigene phylogenetic analyses did not provide finer resolution in the closest living species/genus for the origins of the two plastids (18–22). Such uncertainties are potential drawbacks of cryptophytes and chlorarachniophytes as the model organisms to study the reductive process on the endosymbiont genome during secondary endosymbiosis. Without pinpointing the precise origin of the plastid (or endosymbiotic alga gave rise to the plastid), it is difficult to reconstruct the original gene contents of the nuclear genome of the endosymbionts, and the reductive process that shaped the current nucleomorph genomes in the two algal lineages. Thus, it is ideal to find and investigate a novel nucleomorph-bearing organism, of which endosymbiont origin is resolved at a fine evolutionary scale, and compare with cryptophytes and chlorarachniophytes. However, no novel nucleomorph-bearing lineage has been found since the discovery of the nucleomorph in chlorarachniophytes in 1984 (10).

Dinoflagellates are a eukaryotic group belonging to Alveolata, comprising both photosynthetic and non-photosynthetic species (23, 24). The vast majority of photosynthetic dinoflagellates possess red alga-derived plastids containing a unique carotenoid peridinin (25, 26). It is widely accepted that peridinin-containing plastid has already existed in the ancestral dinoflagellate, and photosynthetic capacity has been lost secondarily on multiple branches of the tree of dinoflagellates (27–29). In addition to multiple losses of photosynthesis, there have been reported multiple, different types of non-canonical plastids lacking peridinin. So far, three types of non-canonical plastids have been known (i) the plastids containing chlorophylls (Chls) *a* and *c* plus 19′ hexanoyloxyfucoxanthin, (ii) those containing Chls *a* and *b* (Chls *a*+*b*), and (iii) those containing Chls *a* and *c* plus fucoxanthin (26, 30–32). The pigment composition together with molecular phylogenies inferred from plastid genes clearly designated the origins of the first, second, and third non-canonical plastids described above as a haptophyte, a green alga, and a diatom, respectively (33–38). In the species with the haptophyte-derived plastids, the endosymbiotic algae are regarded to be fully integrated into the dinoflagellate (host) cells, because no cellular component except the plastid remains, and gene transfer from the endosymbiont genome to the dinoflagellate genome (i.e. endosymbiotic gene transfer or EGT) has been detected (36, 39–45). In *Lepidodinium viride* with the green alga-derived plastid, although a nucleus-like structure in the compartment corresponding to the cytoplasm of the endosymbiont alga was reported (31), it is still controversial (see discussion). On the other hand, the species with the third non-canonical plastids are unique in maintaining major cellular components of the endosymbiont, such as the plastid, nucleus and mitochondrion (46–48). To our knowledge, there has been no molecular evidence for the diatom endosymbionts being modified severely during the endosymbiosis (43, 49–52).

Here, we report two undescribed dinoflagellates, strains MGD and TGD, with green alga-derived plastids containing Chls *a*+*b*. The two dinoflagellates are distinctive from each other in terms of their cell morphologies, and no clear affinity between the two hosts was recovered by molecular phylogenetic analyses. In both MGD and TGD, conspicuous nucleus-like structures with DNA were identified in the periplastidal compartments (PPCs) that correspond to the endosymbiont cytoplasm. We successfully obtained the evidence for the green algal endosymbionts being genetically integrated into the dinoflagellate host cells. Both green algal endosymbionts showed clear phylogenetic affinities to Pedinophyceae, a particular group of green algae. Combining together, we conclude that MGD and TGD are novel nucleomorph-bearing organisms harboring Chls *a*+*b*-containing plastids derived from endosymbiotic Pedinophyceae green algae. Finally, we propose the two dinoflagellates as new experimental models to study secondary endosymbiosis.

## Results

### MGD and TGD possess the nucleomorphs

Cultures of dinoflagellates, strains MGD and TGD were isolated from two distinct coastal locations in Japan. These flagellates commonly contain green-colored plastids, instead of peridinin-containing plastids in major photosynthetic dinoflagellates, but their overall cell structures appeared to be distinct from each other (Figs. 1b and f). The green-colored plastids in the two dinoflagellates are most likely of green algal origin, as pigment profiles chiastic to green algae were detected from MGD and TGD (*SI Appendix*, Fig.S1). Significantly, the cell structure and plastid shape of *Lepidodinium* spp. (53), previously described species bearing green alga-derived plastids, are distinctive from those of MGD and TGD (*SI Appendix*, Fig.S1). Under the transmission electron microscopy (TEM) observations, nucleus-like structures with double-membrane were found in the spaces between the second and third plastid membranes, which corresponds to the cytoplasm of the endosymbiont algae, in MGD and TGD (Figs. 1a, c, e and g). The nucleus-like structures in MGD and TGD are clearly distinct from characteristic dinoflagellate nuclei in the cytoplasm (Figs. 1a and e).

**Figure 1.**
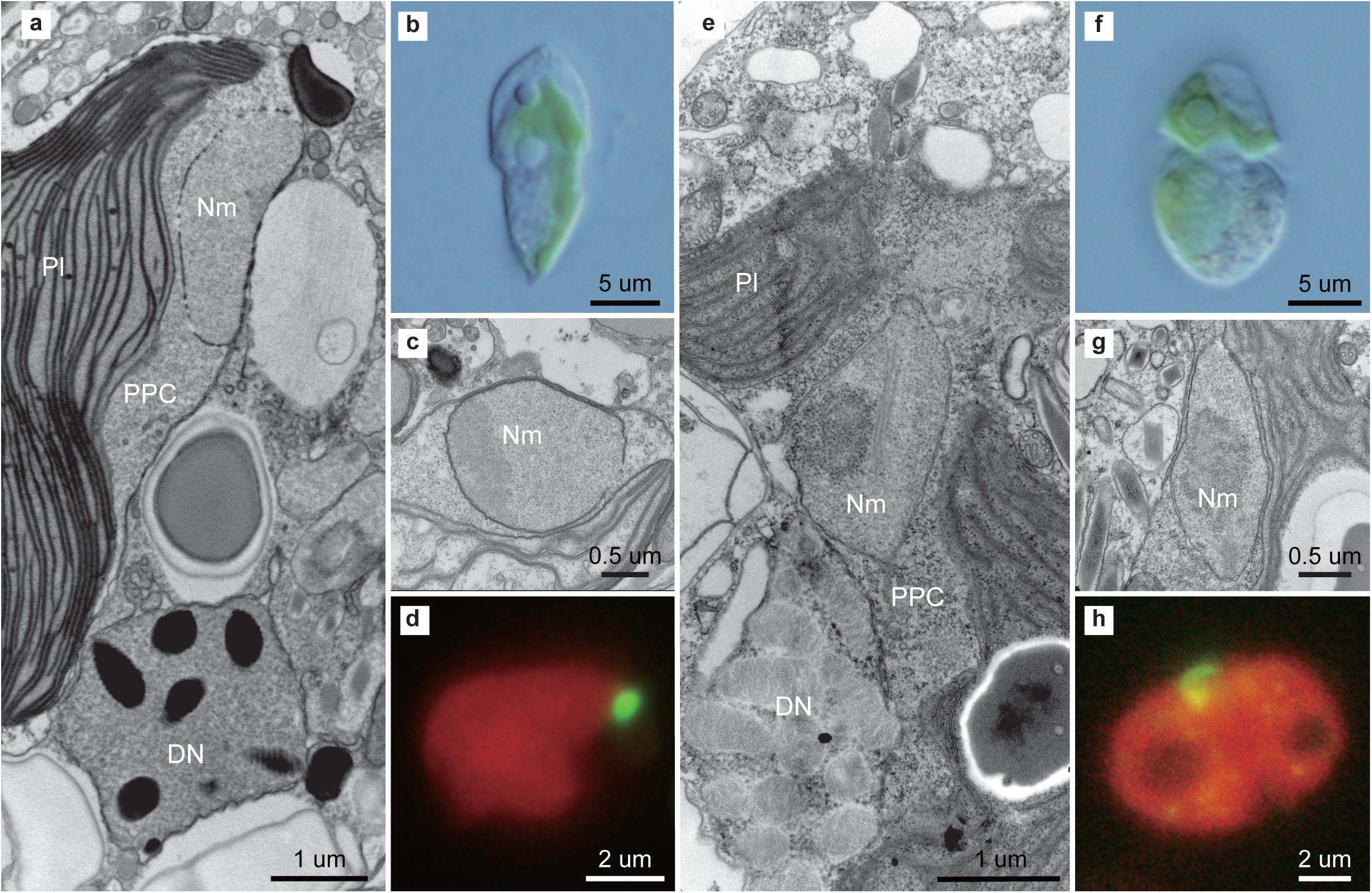
Morphology of undescribed dinoflagellate strains MGD (left) and TGD (right). **a and e** Cross sections of the cell under transmission electron microscopy (TEM), showing the dinoflagellate nucleus (DN), nucleomorph (Nm), plastid (Pl) and periplastidal compartment (PPC). **b and f** Whole cell under light micrographs. **c and g**. Enlargement image of a cross section of the cell under TEM observation. **d and h** Fluorescent microscopy with SYBER green I-staining image.

We conducted fluorescent microscopic observations to examine whether the nucleus-like structure in the PPCs contains any DNA by SYBR green staining. In our preliminary observations staining whole MGD and TGD cells, DNA signal from the endosymbiont compartment was undetectable, due to the over-illumination of the dinoflagellate nucleus (data not shown). Thus, the plastid enclosing the nucleus-like structure was separated from the dinoflagellate nucleus prior to SYBR green staining. We confirmed that the isolated plastids were associated physically with the nucleus-like structures by TEM observation (*SI Appendix*, Fig.S2). The observation of the SYBR green-stained plastid samples successfully detected clear DNA signals on the surface of the plastid, of which locations are consistent with the nucleus-like structures in the PPCs (Figs. 1d and h). The results described above strongly suggest that the nucleus-like structures in the PPCs are derived from the genome-containing nuclei of the green algal endosymbionts in MGD and TGD.

In dinoflagellate species bearing obligate diatom endosymbionts, collectively called ‘dinotoms’ (e.g., *Kryptoperidinium foliaceum* and *Durinskia baltica*), the endosymbionts retain their nuclei and mitochondria as well as the plastids (33, 38). The presence of mitochondria in the endosymbiont alga-derived compartment suggests that energy production in the endosymbiont has yet to be fully under host control in the dinotom system. Thus, the endosymbionts in dinotoms are much less morphologically reduced and host-governed than those in cryptophytes or chlorarachniophytes, which contain the residual nuclei (nucleomorphs) and plastids, but no mitochondrion (7). Our TEM observations detected the ribosomes but no mitochondrion in the PPC of MGD or TGD (Figs. 1a, 1e and S2), suggesting that their green algal endosymbionts were ultrastructurally reduced and governed by the host to the similar level of the endosymbionts in cryptophytes and chlorarachniophytes. The microscopic data described above strongly suggest that (i) the endosymbiont compartments in both MGD and TGD are indeed ‘endosymbiont-derived organelles,’ not obligate endosymbionts, and (ii) the nucleus-like structures found in the PPCs of the two dinoflagellates are equivalent to cryptophyte and chlorarachniophyte nucleomorphs. Thus, we conclude that both MRD and TRD possess the nucleomorphs derived from the nuclei of their green algal endosymbionts.

### Recent genetic integration of the green algal endosymbionts into MGD and TGD

In this section, we provide the evidence for the host and endosymbiont being genetically interlocked to each other in MGD and TGD. From the transcriptomic data of MGD and TGD generated in this study, we predicted 57,983 and 73,589 of transcripts encoding putative proteins, respectively. 543 and 961 out of the putative proteins in MGD and those in TGD, respectively, showed high amino acid sequence similarity to nucleus-encode proteins of free-living green algae. Hereafter, the abovementioned transcripts/proteins are referred to as ‘green algal transcripts/proteins.’

There are two possibilities for which genome (or genomes) the ‘green algal genes’ encoding green algal proteins reside, depending on how the host and endosymbiont interlock each other in the MGD and TGD systems. If the host-endosymbiont interlock at the genetic level has yet to be established in the two systems (as proposed for the dinotom system; Hehenberger et al. 2016) (54), green algal genes are anticipated to be found exclusively in their endosymbiont genomes. Nevertheless, the endosymbiont-derived compartments of MGD and TGD are ultrastructurally more reduced than those of dinotoms, implying that the host-endosymbiont interlock is more advanced in the former systems than the latter. If the host-endosymbiont interlock has reached at the genetic level in the MGD and TGD systems (as observed in cryptophytes and chlorarachniophytes)(12), some of green algal genes were likely transferred from the endosymbiont genome to the host genomes (i.e., endosymbiotic gene transfer or EGT).

We repeated the procedure described above to retrieve the transcripts encoding the proteins conserved among alveolates (including dinoflagellates) in MGD and TGD. Such ‘alveolate transcripts’ were most likely expressed from the host (dinoflagellate) genomes. Alveolate transcripts in MGD formed a cluster in the two-dimensional plot based on the G + C content (GC%) of first codon positions and that of third codon positions (dots in orange; Fig. 2a). Likewise, alveolate transcripts in TGD formed a cluster in the same plot, but shifted toward higher GC% in third codon position (Fig. 2b). In sharp contrast, green algal transcripts from both MGD and TGD were found to be split into two populations (dots in green; Figs. 2a and 2b), and the population with high GC% overlapped with alveolate transcripts. We estimated the abundance of each of green algal/alveolate transcripts in MGD and TGD by calculating FPKM (fragments per kilo-base transcript length per million fragments mapped)(55). Green algal transcripts with high GC% and alveolate transcripts appeared to be expressed at similar levels in both dinoflagellates (*SI Appendix*, Fig.S3). On the other hand, the average FPKM for green algal transcripts with low GC% (334.8 and 279.6 for MGD and TGD, respectively) appeared to be higher than those for the transcripts with high GC% (5.7 and 4.8 for MGD and TGD, respectively). Similar difference in transcriptional intensity between the nuclear and nucleomorph genomes has been documented in cryptophytes and chlorarachniophytes (13, 56).

**Figure 2.**
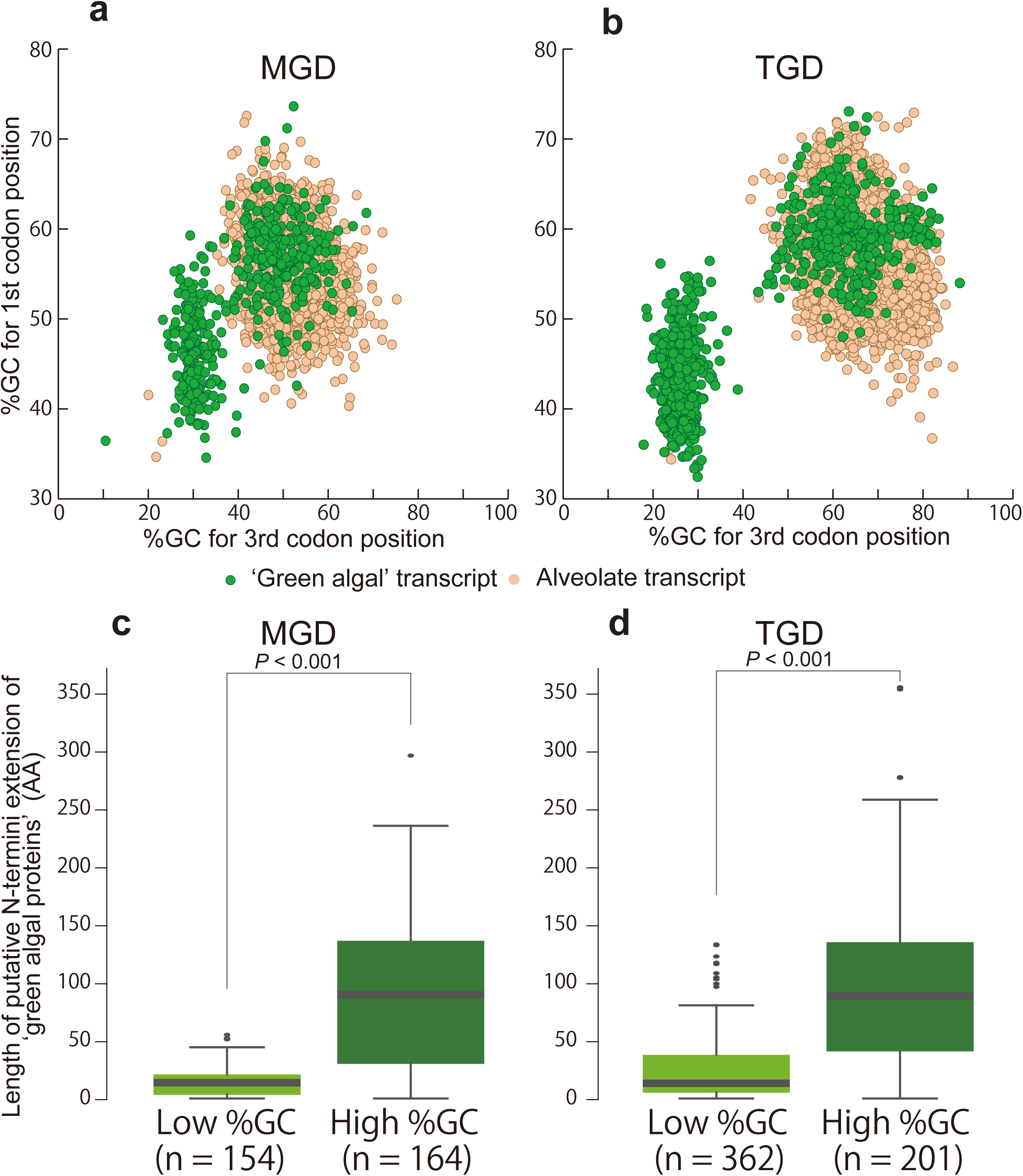
Scatter plots showing the distribution of G + C contents (upper part) and box plots for putative N-terminal extension (lower part) of the transcripts found in TGD (left) and MGD (right). **a and b** X- and Y-axes show the G + C contents (GC%) of first and third codon positions, respectively. Plots in green and orange represent the transcripts encoding the putative green algal and alveolate proteins, respectively. In both plots, green algal transcripts were divided into two populations based on GC%, and the ones with higher GC% overlapped with the masses of alveolate transcripts, which were presumably expressed from the dinoflagellate nuclear genomes. **c and d** Box plots of N-terminal extension of ‘green algal’ transcripts with low and high GC%. X-axes show lengths of putative N-terminal extensions estimated based on comparison with homologs of free-living green algae (see Material and Methods). *P* values displayed in the plots were calculated based on the Wilcoxon rank-sum test.

Mature mRNA molecules transcribed from dinoflagellate nuclear genomes are known to possess a particular short sequence (spliced leader or SL sequence) at their 5′ termini (57). Thus, green algal transcripts expressed from the host nuclear genome are anticipated to be preceded by the SL sequences. We examined the presence/absence of the SL sequence in six green algal transcripts identified from TGD by RT-PCR using a forward primer matching the dinoflagellate SL sequence (57) and reverse primers matching specifically to the individual transcripts. As presented in *SI Appendix*, Fig.S 4, the 5′ termini of the three transcripts with high GC% were successfully amplified, while no amplification was observed for the three transcripts with low GC% (Upper panel in *SI Appendix*, Fig.S4). We subjected six green algal transcripts identified from MGD to the same RT-PCR experiments, and observed the amplification of the 5′ termini occurred only for the transcripts with high GC% (Lower panel in *SI Appendix*, Fig.S4). These transcripts bearing the SL sequence are the compelling evidence that the host genomes of MGD and TGD carry and transcribe the genes acquired from the genomes of their green algal endosymbionts. Although we experimentally examined the presence/absence of the SL-sequence in a subset of green algal transcripts (see above), we predict that the majority of those with high GC% were expressed from the dinoflagellate nuclear genomes and received the SL sequences post-transcriptionally in both MGD and TGD. We here conclude that the host and endosymbiont in both MGD and TGD as the partner interlocked to each other by EGT, as those in cryptophytes and chlorarachniophytes.

Green algal transcripts with low GC% (Figs. 2a and 2b) likely bear no SL sequence at their 5′ termini, implying the presence of the second genomes, of which GC% are lower than the dinoflagellate nuclear genomes, in both MGD and TGD. Accumulated genomic data clearly suggest that endosymbiont and organelle genomes underwent reductive evolution (e.g., cryptophyte and chlorarachniophyte nucleomorph genomes) tend to bear low GC% (7). Considering the reduced characteristics in the endosymbiont compartments in MGD and TGD at the morphological level, we predict that the nucleomorph genomes in the two dinoflagellates bear lower GC% than those of the dinoflagellate nuclear genomes. Thus, the sources of green algal transcripts with low GC% are most likely the nucleomorph genomes. It is worthy to note that ‘house-keeping’ proteins, which are involved in the eukaryote-type machineries for translation, transcription and replication, were found to be encoded almost exclusively by green algal transcripts with low GC% (i.e. putative nucleomorph transcripts) in both MGD and TGD (*SI Appendix*, Fig.S5a and b). In contrast, both transcripts encoding proteins involved in plastid metabolic pathways appeared to span the two populations of green algal transcripts with distinct GC% (*SI Appendix*, Fig.S5c and d). Similar biased gene content in the nucleomorph genomes has been documented in the cryptophyte and chlorarachniophyte systems (7, 12).

We identified the green algal transcripts with high GC% (i.e. putative nuclear transcripts) encoding ‘plastid-related proteins’ involved in plastid function and maintenance (e.g., photosynthesis) in both MGD and TGD (*SI Appendix*, Fig.S5c and d). In theory, to operate the plastids, these nucleus-encoded plastid-related proteins (many of those are presumably acquired from the green algal endosymbionts) need to be targeted into the plastids after being synthesized by the host machinery in the cytoplasm in MGD and TGD.

In photosynthetic eukaryotes (e.g., green algae) with primary plastids surrounded by two membranes, many of nucleus-encoded plastid-targeted proteins bear N-terminal extensions (so-called ‘transit peptides’ or ‘TPs’) that work as plastid-targeting signal. The N-terminal extensions of the nucleus-encoded proteins targeted into complex plastids, which are surrounded by three or four membranes, tend to have a bipartite structure comprising a hydrophobic ‘signal peptide (SP)’ followed by the TP-like (TPL) region (58). As both MGD and TGD plastids are surrounded by four membranes (Fig. 1a and 1e), the nucleus-encoded plastid-targeted proteins of the two dinoflagellates should have the bipartite plastid-targeting signal. Nevertheless, the proteins encoded by endosymbiotically transferred green algal genes unlikely equipped the bipartite signal *ab initio*, and need to have modified their N-terminal extensions by appending the SPs to be targeted into the current MGD and TGD plastids. If the above scenario is the case, the nucleus-encoded plastid-targeted green algal proteins in MGD and TGD possess the N-terminal extensions longer than those of the homologous proteins in free-living green algae with primary plastids.

It is reasonable to expect that the nucleus-encoded plastid-targeted green algal proteins are a subset of the green algal proteins encoded by the putative nuclear transcripts (i.e. green algal transcripts with high GC%). As expected, the nucleus-encoded green algal proteins tend to bear apparently longer N-terminal extensions than the counterparts in free-living green algae (Fig. 2c and d), implying that these N-terminal extensions have bipartite structures. Indeed, some green algal proteins in MGD and TGD were predicted to have the typical bipartite plastid-targeting signals (*SI Appendix*, Fig.S6). The results described above suggest that both MGD and TGD possess the nucleus-encoded green algal proteins, which are localized post-translationally in the plastid. On the other hand, the green algal proteins encoded by green algal transcripts with low GC% (i.e. putative nucleomorph transcripts) are unnecessary to have the N-terminal extensions with the bipartite structure. The difference in length of the N-terminal extensions between the nucleomorph-encoded green algal proteins and the counterparts in free-living green algae was less obvious than the first comparison (Fig. 2c and 2d).

Our detailed assessment focused on green algal transcripts identified multiple *psbO* and *rbcS* transcripts with distinct GC% in MGD (Fig.3). Those green algal transcripts showed high affinity to particular green algae, *Pedinomonas* (See the next section for the details). We confirmed the SL sequences of the high-GC% versions of the aforementioned transcripts by bioinformatically or experimentally (see above), but yielded no evidence for the SL sequence at the 5′ termini of the low-GC% counterparts (Fig. 3). These data suggest that MGD possesses two sets of *psbO* and *rbcS* genes, one is nucleus-encoded (high-GC% and generate the SL-bearing transcript) and the other is nucleomorph-encoded (low-GC% and generate the SL-lacking transcript). Likewise, TGD likely possesses both nucleus- and nucleomorph-encoded *petC* genes, as we found two *petC* transcripts, one is of high-GC% and bears the SL sequence, and the other is of low GC% and bears no SL sequence.

**Figure 3.**
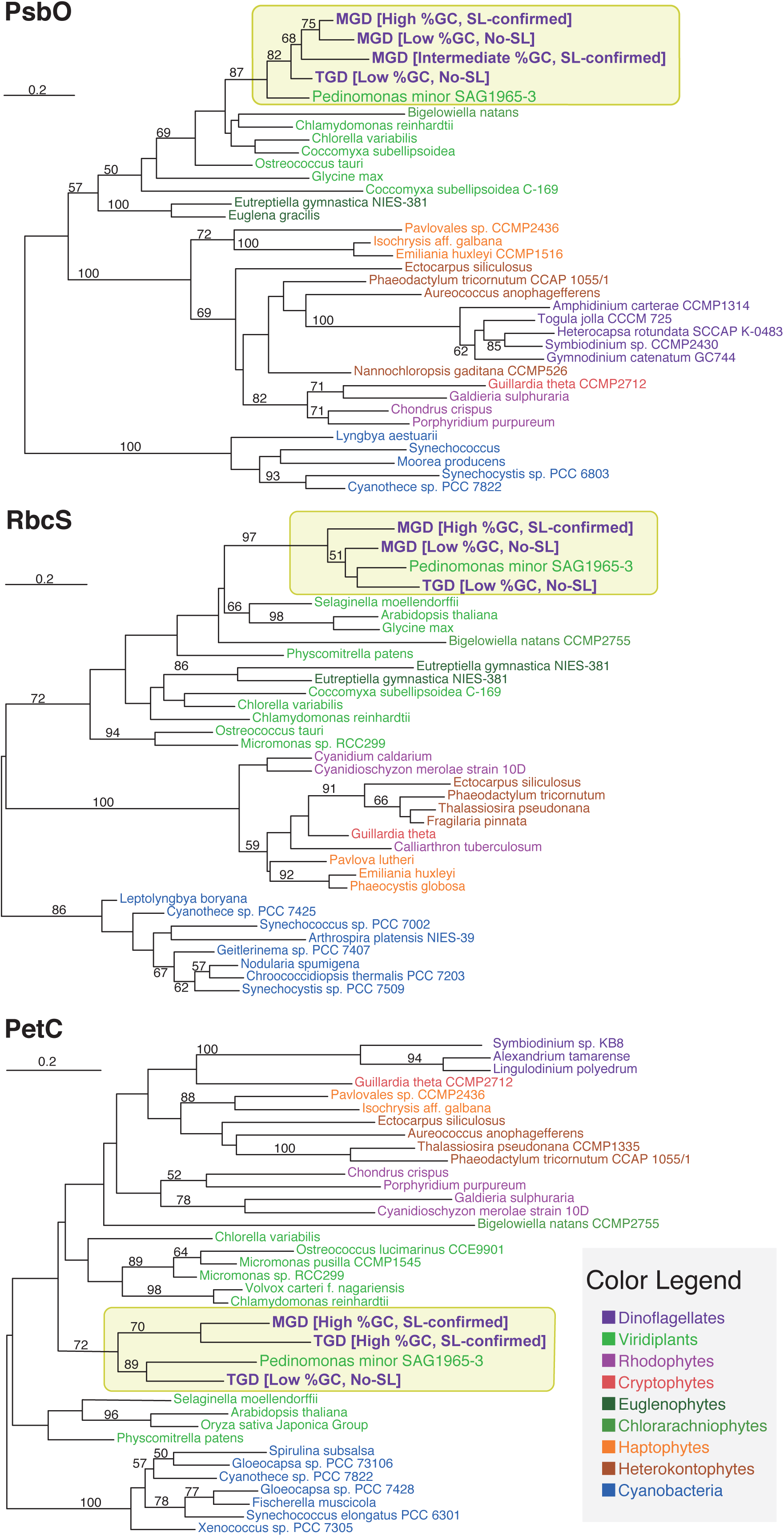
Maximum likelihood (ML) trees for the green algal orthologous proteins with distinct GC%. The numbers above branch show non-parametric ML bootstrap values. Only ML bootstrap support values greater than 50% are shown on the corresponding branches.

In addition to the nuclear (high-GC%) and nucleomorph (low-GC%) versions described above, we found the third *psbO* transcript, of which GC% cannot be defined as low or high, in MGD (Fig. 3). The *psbO* transcript with an ‘intermediate GC%’ was found to bear the SL sequence, indicating that MGD possesses a nucleus-encoded *psbO* gene, of which GC% does not match to other nuclear genes including the high-GC% version of *psbO* gene. We here propose that the intermediate-GC% version of *psbO* gene was transplanted in the nuclear genome before the GC% of the nucleomorph genome had been lowered at the current level. Alternatively, the intermediate-GC% *psbO* gene is the result of more recent EGT than the high GC% one, and the former gene has not accommodated its GC% to those of other nucleus-encoded genes.

### Pedinophyceae origin of the green algal endosymbionts in MGD and TGD

We isolated two eukaryotic small subunit rRNA (18S rRNA) genes—one with and the other without a clear phylogenetic affinity to those of dinoflagellates—from each of MGD and TGD. The 18S rRNA sequences isolated from MGD and TGD were phylogenetically analyzed along with those from red algae, green plants including green algae and land plants, glaucophytes, cryptophytes, chlorarachniophytes, and dinoflagellates. The 18S rRNA phylogeny placed ‘non-dinoflagellate type’ MGD and TGD sequences within the sequences from free-living green algae, being distant from the clade of the dinoflagellate sequences including ‘dinoflagellate type’ MGD and TGD sequences (Fig. 4). We thus regard the positions of dinoflagellate type and non-dinoflagellate type sequences from MGD and TGD in the 18S rRNA phylogeny as the phylogenetic positions of the host and endosymbiont in each of the MGD and TGD systems.

**Figure 4.**
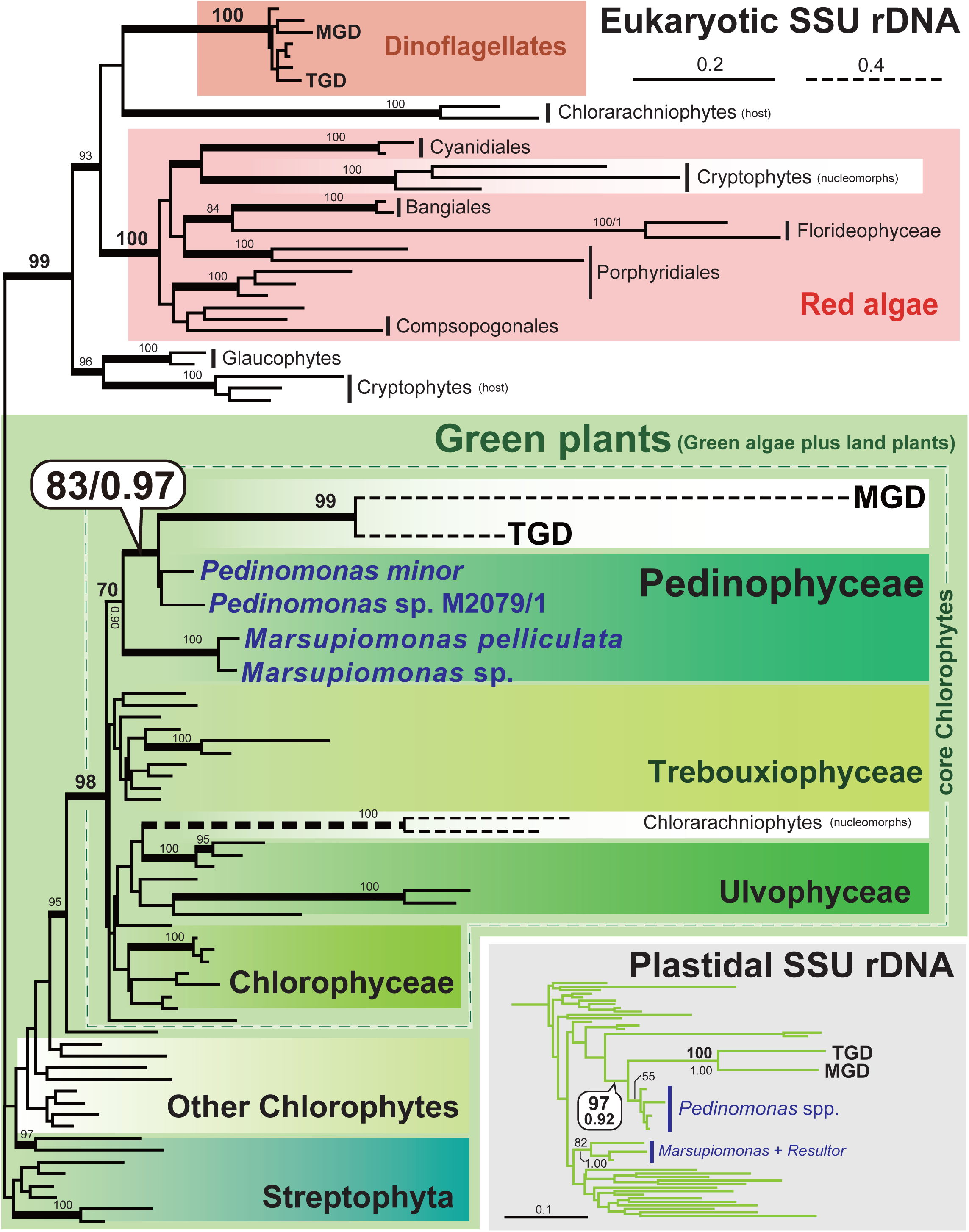
Maximum likelihood (ML) tree inferred from eukaryotic small subunit ribosomal RNA (18S rRNA) sequences. All the taxon names are omitted except MGD, TGD, and Pedinophyceae green algae. Taxon labels of red and green algae are followed Adl et al. (2019)(75). Only ML bootstrap support values greater than 80% are shown on the corresponding branches. The branches supported by Bayesian posterior probabilities greater than 0.95 are shown as thick lines. The clade comprising MGD, TGD, and Pedinophyceae green algae inferred from plastidal small subunit rRNA (16S rRNA) sequences is shown in the box. The 16S rRNA tree including *Lepidodinium chlorophorum* with full taxon names is provided as *SI Appendix*, Fig.S7.

Unlike cryptophyte or chlorarachniophyte nucleomorphs, the 18S rRNA phylogeny clarified the precise origins of the green algal endosymbionts in the MGD and TGD systems. Non-dinoflagellate type MGD and TGD sequences grouped together, and this MGD-TGD clade was placed within a clade of a small collection of green algae, Pedinophyceae, with the specific affinity to *Pedinomonas* spp. (Fig. 4). The monophyly of *Pedinomonas* plus MGD and TGD received a maximum-likelihood bootstrap value (MLBP) of 83 % and a Bayesian posterior probability (BPP) of 0.97 (Fig. 4). The detailed origins of the endosymbionts in MGD and TGD were also assessed by phylogenetic analyses of plastidal small subunit rRNA (16S rRNA) sequences (Fig. 4). The plastidal 16S rRNA phylogeny united MGD, TGD, and *Pedinomonas* spp. into a clade with a MLBP of 97% and a BPP of 0.92, excluding other Pedinophyceae considered in the analyses, *Marsupiomonas* spp. and *Resultor mikron* (Fig. 4). The two phylogenetic analyses consistently and strongly indicate that the current plastids in both MGD and TGD were traced back to Pedinophyceae green algae closely related to *Pedinomonas*.

Some of us have already reported the Pedinophyceae origin of the Chls *a*+*b*-containing plastids in the dinoflagellate genus *Lepidodinium* (59). Indeed, in the plastidal 16S rRNA phylogeny including the *L. chlorophorum* sequence, *L. chlorophorum* robustly grouped with MGD and TGD, and this dinoflagellate clade as a whole was found to be sister to *Pedinomonas* (*SI Appendix*, Fig.S7; Note that the *L. chlorophorum* sequence was excluded from the analyses presented in main Figure 4 due to its extremely divergent nature). The simplest interpretation of this tree topology is that *L. chlorophorum*, MGD, and TGD share a single ancestor with a Pedinophyceae-derived plastid. Nevertheless, a taxon-rich dinoflagellate phylogeny inferred from eukaryotic large subunit rRNA (28S rRNA) sequences (Fig. 5 and *SI Appendix*, Fig.S8) appeared to be inconsistent with the scenario deduced from the plastid phylogeny. In the host phylogeny deduced from 28S rRNA sequences, MGD or TGD showed no particular affinity to any other dinoflagellates, while *Lepidodinium* was nested within a robustly supported clade of dinoflagellates with peridinin-containing plastids (e.g., *Gymnodinium* and *Nematodinium*). We examined the relationship among *Lepidodinium*, MGD, and TGD by subjecting the ML and four alternative trees to an approximately unbiased test (60). In the alternative trees, all or subsets of the three species were enforced to be monophyletic (Table 1). As results, Tree nos. 2, 3, and 5, in which *Lepidodinium* was connected to either MGD or TGD, or both, were rejected with very small *p* values, while only Tree no. 4 bearing the monophyly of MGD and TGD failed to be rejected (Table 1). Thus, we can conclude that the host lineages of the three species are highly unlikely be monophyletic.

**Figure 5.**
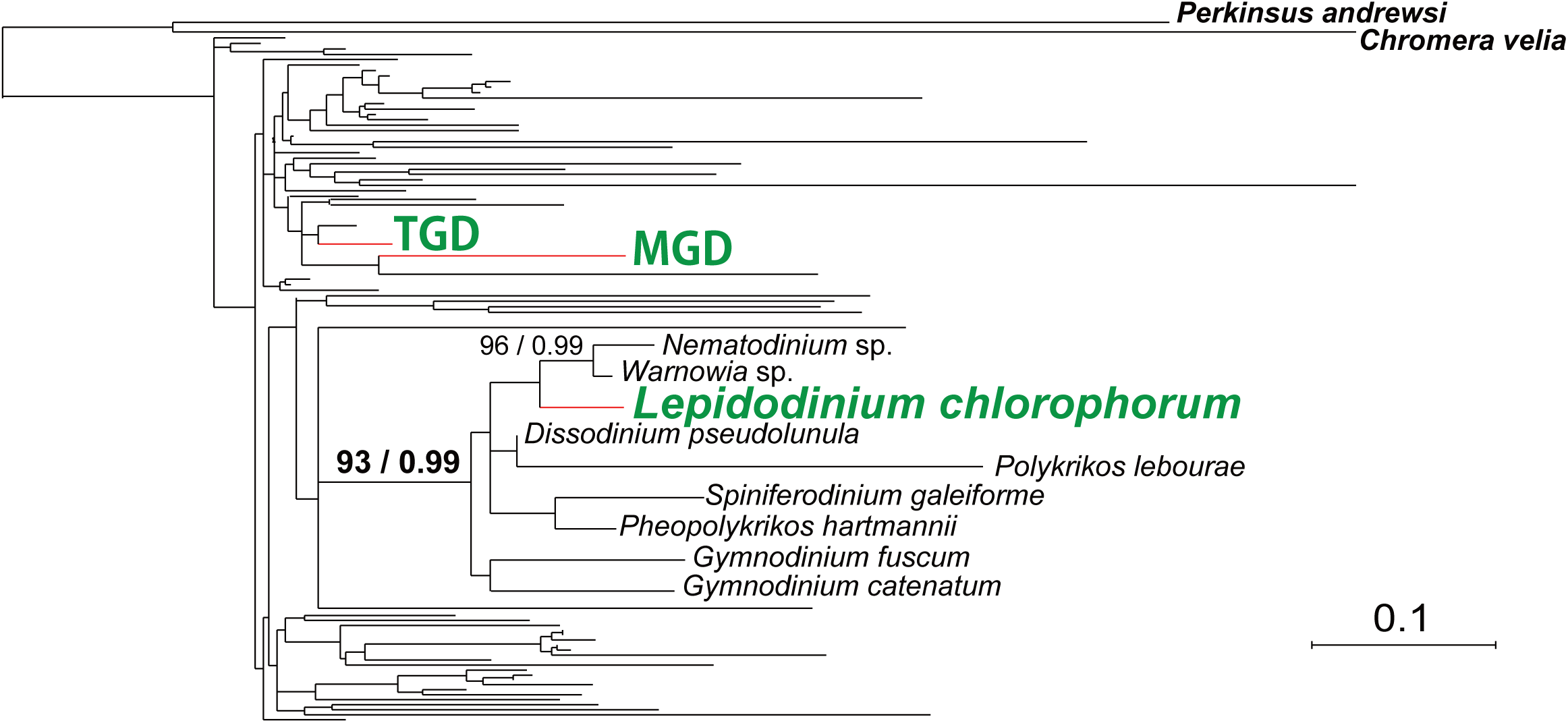
Maximum likelihood (ML) tree inferred from eukaryotic large subunit ribosomal RNA (28S rRNA) sequences. The topology in question was emphasized. Only ML bootstrap support values greater than 80% are shown on the corresponding branches. The branches supported by Bayesian posterior probabilities greater than 0.95 are shown. The same tree with full taxon names is shown in *SI Appendix*, Fig.S8.

**Table 1.** Approximately unbiased tests for phylogenetic relationships amongst *Lepidodinium*, MGD, and TGD

| Nos | Constraints | $\Delta\ln Ls$ | *p* values |
| --- | --- | --- | --- |
| 1 | - (ML tree) | - | 0.736 |
| 2<sup>a</sup> | Monophyly of *Lepidodinium*, MGD, and TGD | 115.2 | 4e-06* |
| 3<sup>a</sup> | Monophyly of *Lepidodinium* and MGD | 64.4 | 0.002* |
| 4<sup>a</sup> | Monophyly of MGD and TGD | 8.8 | 0.285 |
| 5<sup>a</sup> | Monophyly of *Lepidodinium* and TGD | 115.8 | 1e-04* |

\*: rejected by 1% criterion.

The inconsistence between the host and plastid phylogenies demands the Pedinophyceae endosymbiosis in the ancestral *Lepidodinium* species to be separated from that (or those) gave rise to the current plastids in MGD and TGD. Furthermore, MGD and TGD plastids may have been established through separate Pedinophyceae endosymbioses after the divergence of dinoflagellates, as no intimate affinity between MGD and TGD was recovered in the 28S rRNA phylogeny representing the host relationship (Fig. 5 and *SI Appendix*, Fig.S8).

We currently cannot rationalize why separate dinoflagellate lineages engulfed very closely related Pedinophyceae green algae as endosymbionts. To answer the above question, we need to understand the interaction between dinoflagellates and Pedinophyceae at the genetic, physiological, and environmental levels.

## Discussion

In this study, we reported previously undescribed dinoflagellates, strains MGD and TGD, both of which still retain the remnant intercellular structures of algal endosymbionts enclosing the nuclei and plastid, but no mitochondrion (Fig. 1). Multiple cases of EGT in MGD and TGD indicate that the endosymbiont-derived compartments had been integrated as the organelles into the two dinoflagellates. Combining these aspects, we regard MGD and TGD as novel nucleomorph-bearing organisms. Cryptophytes and chlorarachniophytes, have been investigated intensively as model organisms to depict organellogenesis that transformed an algal endosymbiont into the plastid (e.g., Curtis et al. (2012), Tanifuji and Onodera (2017))(12, 61). Curiously, three characteristics in the cryptophyte and chlorarachniophyte nucleomorphs were found in those of MGD and TGD (see below). Pioneering studies demonstrated that, in both cryptophytes and chlorarachniophytes, (i) the nucleomorph genomes are rich in house-keeping genes, (ii) the G + C% of the nucleomorph genomes are low, as seen in other reduced genomes such as plastid and mitochondrial genomes and (iii) nucleomorph genes tend to be transcribed more intensively than nuclear genes 61. Although no genome data is available for the nucleomorphs of MGD or TGD, the putative nucleomorph transcripts of the two dinoflagellates appeared to be rich in those encoding house-keeping proteins (*SI Appendix*, Fig.S5). We also noticed that nucleomorph genes were found to be transcribed more intensively than nuclear genes in both MGD and TGD (*SI Appendix*, Fig.S3). We suspect that the two characteristics in gene content and transcription shared among the nucleomorph genomes known to date might provide the critical clues to understand organellogenesis.

Besides the characteristics shared with cryptophytes and chlorarachniophytes (see above), MGD and TGD turned out to equip characteristics that are not available in the two previously known nucleomorph-containing lineages. The phylogenetic analyses inferred from plastidal 16S and eukaryotic 18S rRNA sequences revealed that the plastid origins of MGD, TGD and *Lepidodinium* spp. are the close relatives of a particular green algal genus, *Pedinomonas* (Fig. 4). In contrast, neither origin of the red alga engulfed by the ancestral cryptophyte nor that of the green alga engulfed by the ancestral chlorarachniophyte has been pinpointed at the genus level (19–22). Thus, the modifications on the endosymbiont genomes during the transition from an endosymbiotic alga into the plastid can be predicted directly by comparing MGD/TGD nucleomorph genomes to those of free-living Pedinophyceae in the future.

The process transferred endosymbiont genes to the host genome may have gone through a transitional state in which the particular genes co-occurred in both endosymbiont and host genomes 13, 56. Nevertheless, Curtis et al (2012) (12) proposed that both cryptophyte and chlorarachniophyte systems are in the post-EGT state, as little evidence for DNA flux from the nucleomorph to nuclear genomes was found in the two systems. In contrast, we identified both nuclear and nucleomorph versions of *psbO* and *rbcS* genes in MGD, and those of *petC* gene in TGD (Fig. 3). Moreover, the MGD nuclear genome appeared to possess two *psbO* genes with distinct GC% (Fig. 3), which are likely the outcome from two separate EGT events. The gene redundancy between the nuclear and nucleomorph genomes in MGD and TGD suggest that EGT has yet to be competed in either of the two dinoflagellate systems. In this sense, MGD and TGD are excellent models to elucidate the detailed process of EGT during organellogenesis.

Jackson et al. (2018) proposed that the current plastids in *Lepidodinium* spp. were established more recently than the chlorarachniophyte plastids (62). Both *Lepidodinium* plastids and MGD/TGD plastids were derived from closely related Pedinophyceae green algae (Fig. 4), but the host phylogeny inferred from eukaryotic 28S rRNA sequences clearly suggest that separate endosymbioses gave rise to the plastids in MGD and TGD, and those in *Lepidodinium* spp. (Fig. 5 and Table 1). Although the nucleus-like structure within plastid in *Lepidodinium viride* was reported previously (31), at least clear structure such shown in MGD and TGD were not observed in our survey. The difference in intracellular structure between MGD/TGD and *Lepidodinium* spp. leads us to propose that the endosymbiosis in MGD/TGD was more recently than that in the ancestral *Lepidodinium* cell (or that in the ancestral chlorarachniophyte cell). We suspect that (i) the clarity in plastid/nucleomorph origin and (ii) traits correspond to the transitional state of EGT (see above) found in MGD and TGD stem from their ‘evolutionary youthfulness.’

The current study reports the third nucleomorph-bearing organisms, dinoflagellate strains MGD and TGD, which were discovered the first time in last 30 years, and anticipated to shed a light the nature of endosymbiogenesis. The two dinoflagellates will be the foundation of future works that deepen our knowledge from the pioneering works on cryptophytes and chlorarachniophytes, which were used to be the sole models to study the evolutionary process transforming an algal endosymbiont into the plastid.

## Materials and methods

### Strains and culture condition

Two novel green dinoflagellate strains MGD and TGD were collected from coastal areas in Muroran, Hokkaido, Japan and Tsuruoka, Yamagata, Japan in September 2011, respectively. Single cells were isolated using a glass pipette under the light microscopy. The strains were cultivated in 1/2 final concentration of Daigo IMK medium (Wako, Osaka, Japan) with seawater at 20 °C under a light/dark cycle of 12/12 h.

### Light and fluorescent microscopy

Living cells were observed by Olympus IX71 inverted microscope (Olympus, Tokyo, Japan) with Olympus DP71 CCD camera (Olympus). For fluorescent microscopic observation, the cells were fixed with the same volumes of fixation buffer containing 2.5% glutaraldehyde, 2% paraformaldehyde, 0.2M sucrose and 0.1M cacodylate at pH 7.0 for 5 min at room temperature. Fixed cells were rinsed three times by distilled water in 10 min at room temperature, and were harvested by centrifugation at 2,810 *g* for 10 min at 18 °C. The cells were put onto a glass coverslip coated by poly-L-Lysin (Wako), and left to stand for 30 min. DNAs in the fixed cells were stained by 0.1% SYBR green I solution for 10 min at room temperature. After washing three times with distilled water for 10 min, the DNA-stained cells were mounted with ProLong Diamond Antifade Mountant (Life Technologies), and then observed using Leica DMRD light and fluorescence microscope (Leica, Wetzlar, Germany) with an Olympus DP73 CCD camera (Olympus).

### Transmission electron microscopy

The first fixation step was carried out as the same method described above, then, the cells were washed with 0.2M cacodylate buffer. The second fixation was performed with buffer containing 1% OsO4 and 0.2M cacodylate for 15 min at 4 °C. Cells were dehydrated by various concentrations of ethanol series in 10 min for each, and embedded in low viscosity resin via polypropylene oxide three times in one hour at room temperature. Embedded samples were polymerized for 14 hours at 70 °C. Polymerized block was sectioned using a diamond knife and placed onto formvar-coated copper grids. Ultra-thin sections of the cells were stained with 2% uranyl acetate and lead citrate. Observations were carried out using H7650 (Hitachi, Tokyo, Japan).

### Plastid isolation

Living TGD cells were centrifuged for 10 min at 2,810 *g* to make the cells burst by physical pressure of centrifugation. For MGD, the cells were put into 25 % final concentration of sucrose solution for 5 min. Then, 4 volumes of distilled water were added to make the cells burst by hypoosmotic shock. TGD and MGD cell pellets including the plastids freed from other cellular structures were harvested by centrifugation, and were fixed as described above. The freed plastids were sought and taken photos under the fluorescent microscopy and the transmission electron microscopy.

### Transcriptome analyses and protein prediction

Total RNAs of MGD and TGD cells were extracted using TRIzol® reagent (Life Technologies). After enrichment of polyA-tailed mRNA molecules, RNA samples were reverse-transcribed and the cDNAs were ligated with adaptor primers. The cDNA libraries from MGD and TGD were then sequenced by HiSeq 2000 instrument (Illumina). From the analyses, 286 and 411 million reads (paired-end) were generated from MGD and TGD libraries, respectively. Sequence reads with low sequencing quality were removed using FASTQ Trimmer and FASTQ Quality Filter programs included in the FASTX-toolkit package (http://hannonlab.cshl.edu/fastx_toolkit/). The remaining reads of MGD and TGD were then assembled into 286,124 and 393,513 transcript contigs using the TRINITY package (release 2013-02-25) (63, 64), respectively. All the transcript contigs were subjected to blastx analysis against an in-house database of protein sequences retrieved from phylogenetically diverse organisms. In case of open reading frames (ORFs) encoding known proteins being identified by blastx analyses, they were translated into amino acid sequences. Otherwise, the longest possible ORFs seen in individual transcripts were translated into amino acid sequences. The putative proteins predicted from the transcriptomic data were further analyzed as described below. All transcriptome data are available from the DNA Data Bank of Japan under BioProject PRJDB8237.

### Functional annotation of the proteins predicted from MGD and TGD transcriptomic data

The proteins predicted from MGD and TGD transcripts were roughly annotated by referring to Kyoto Encyclopedia of Genes and Genomes (KEGG) database (65). First, total 193,301 proteins with KEGG orthology IDs (KO IDs) from 40 eukaryotic and 32 bacterial species were retrieved from KEGG database. The protein sets predicted from each of MGD and TGD were subjected to blastp analyses against retrieved KEGG proteins. Then the eukaryotic sequences retrieved from KEGG database subjected to blastp analyses in the opposite direction (i.e., blastp search against the protein sets predicted from the MGD and TGD transcriptomes). If a particular MGD/TGD sequence and a KO ID matched in reciprocal blastp analyses, the KO ID was assigned to the sequence of interest. We assigned functional annotations to 57,983 and 73,589 of the putative proteins identified from MGD and TGD transcriptome data, respectively.

### Evolutionary origins of the proteins predicted from MGD and TGD transcriptomic data

The phylogenetic origins of functionally annotated proteins were individually assessed as described below. Each of MGD and TGD proteins with functional annotations was subjected to blastp analyses against a custom protein database containing the genome-wide protein data from 129 phylogenetically diverged organisms (48 eukaryotic, 68 bacterial and 13 archaeal species), and the proteins encoded in the plastid genome of a Pedinophyceae green alga, *Pedinomonas minor* (66), a free-living relative of the green algal endosymbionts engulfed by MGD and TGD (see the main text). We identified two sets of proteins, “green algal proteins” and “alveolate proteins,” of which the top five hits from blastp searches were occupied by sequences from members of Viridiplantae or those from alveolates, respectively. MGD and TGD proteins, which hit those encoded in the *P. minor* plastid genome within the top five hits in blastp analyses, were designated as plastid genome-encoded proteins, and were not analyzed any further.

### GC-content and FPKM calculation for each transcript

GC-content for an entire sequence as well as for first, second and third codon positions of each transcript was calculated using an in-house Ruby script. In this study, the green algal transcripts with >50% GC at 1st and >40% GC at 3rd codon positions were designed as high-GC% green algal transcripts, and those with <60% at 1st and <40% at 3rd codon positions were designed as low-GC% green algal transcripts.

To estimate expression levels for each transcript from MGD and TGD, we calculate relative abundances as FPKM (fragments per kilo-base transcript length per million fragments mapped). To calculate FPKM, RNA-seq reads from MGD and TGD were separately aligned onto respective transcriptome assembly using bowtie (67). FPKM were then calculated using RSEM (68) based on the short-read alignments. We carried out those steps using a Perl script (align_and_estimate_abundance.pl) bundled with Trinity (ver. 2.0.6)(63, 64, 68).

### Estimation of N-terminal extensions in green algal proteins

Possible N-terminal extensions of the “green algal proteins” form MGD and TGD were determined based on blastp alignments. First, all of the “green algal proteins” from the two dinoflagellates were subjected to a blastp search against the custom protein database described above. For each “green algal protein,” we defined the mature protein region, which are conserved with top 10 hits from the blastp search. The N-terminus of the putative mature protein region for each protein was set as an average of the start positions of top 10 blast alignments (outliers detected by the Grubbs’ test were not used in the average calculation). In this way, the putative N-terminal extension of each protein corresponds to the amino-acid sequence between the first methionine of the protein sequence and the estimated N-terminus of mature protein region. If no methionine was found in the upstream of a putative mature protein region, we regarded that the particular transcript most likely encodes an N-terminus truncated protein and excluded from the analysis described below. Finally, lengths of putative N-terminal extensions between “green algal proteins” encoded by high-GC% transcripts and those encoded by low-GC% transcripts (see above), were compared. Difference in the length of N-terminal extension between the two groups of “green algal proteins” was tested by the Wilcoxon rank-sum test after removal of the outliers determined by the Grubbs’ test.

### Confirmation of the spliced leader (SL) sequence at the 5′ termini of MGD and TGD transcripts

Total RNA samples were extracted from the cultured MGD and TGD cells using TRIzol® reagent (Life Technologies). Reverse-transcription and cDNA amplification were performed with SMARTer® PCR cDNA Synthesis Kit (Clontech Laboratories, Inc.) according to the manufacturer’s instructions. DNA amplification was carried out as described below. The first PCR was performed with the cDNA sample as the template and a set of primers, an adaptor primer supplied in the kit mentioned above (5’ PCR Primer II A) and a transcript-specific primer. The thermal cycle was set as: 5 cycles consisting of 10 sec at 98°C and 1 min at 68°C; 5 cycles consisting of 10 sec at 98°C, 20 sec at 60°C, and 1 min at 68°C followed by 25 cycles consisting of 10 sec at 98°C, 20 sec at 53°C, 1 min at 68°C. We conducted the second PCR with a SL sequence specific primer 57 and another transcript-specific primer that was set in the nested position to the first primer. The amplicons of the first PCR were used as the template in the second PCR. The thermal cycle was set as: 5 cycles consisting of 10 sec at 98°C and 1 min at 68°C; 5 cycles consisting of 10 sec at 98°C, 20 sec at 60°C, and 1 min at 68°C followed by 25 cycles consisting of 10 sec at 98°C, 20 sec at 53°C, 1 min at 68°C.

### Phylogenetic analyses of ribosomal RNA (rRNA) genes

Eukaryotic small and large subunits rRNAs (18S and 28S rRNA, respectively), and plastidal small subunit ribosomal RNA (16S rRNA) sequences were amplified by the standard PCR and then sequenced by Sanger method. The determined sequences were separately aligned by the program MAFFT (69). After manual exclusion of ambiguously aligned positions, 1,658 position and 82 taxa remained in the final 18S rRNA alignment, 732 positions and 78 taxa in the 28S rRNA alignment, and 1,232 positions and 66 taxa in the plastidal 16S rRNA alignment. The maximum-likelihood (ML) phylogenetic analyses of the three alignments were constructed using RAxML ver. 8.0.2 (70) under the GTR substitution model incorporating among-site rate variation approximated by a discrete gamma distribution with four categories (For the 18S rRNA analysis, the proportion of invariant sites was also incorporated to approximate among-site rate variation). The ML tree was heuristically searched from ten distinct maximum-parsimony (MP) trees, each of which was reconstructed by random stepwise addition of sequences. One thousand bootstrap replicates were generated from the 18S rRNA alignment, and subjected with the rapid bootstrap method under the CAT model using RAxML. For the bootstrap analyses of both 28S rRNA and plastid 16S rRNA alignments, 100 replicates were generated, and individually subjected to tree search initiated from a single MP tree reconstructed by random stepwise addition of sequences using RAxML.

The three alignments were also analyzed by Bayesian method using the CAT-GTR + Γ model implemented in the program PHYLOBAYES v3.3 (71) with two independent chains. Markov chain Monte Carlo chains (MCMC) were run for 80,000 (18S rRNA), 80,000 (28S rRNA) and 78,000 (plastidal 16S rRNA) generations with burn-in of 20,000 generations, respectively. We regarded the two chains for both analyses were converged, as the maxdiff values became less than 0.1. After “burn-in,” the consensus tree with branch lengths and Bayesian posterior probabilities (BPPs) were calculated from the rest of the sampled trees.

### Phylogenetic analyses of overlap green algal protein

Homologous sequences of three green algal proteins in MGD and TGD, namely PsbO, RbcS and PetC, were retrieved from an in-house database comprising proteins sequences from phylogenetically diverse organisms by similarity search using blastp software. Retrieved protein sequences and the homologous sequences from MGD and TGD were then aligned by MAFFT (69) with the L-INS-I method. Ambiguously aligned positions and gaps were removed from the final PsbO, RbcS and PetC alignments, which comprise 35 sequences with 254 amino-acid positions, 35 sequences with 100 amino-acid positions and 36 sequences with 178 amino-acid positions, respectively. Maximum likelihood trees were inferred from the three alignments by IQ-TREE (72) with non-parametric bootstrapping (100 replicates) under the LG4X substitution model.

### Approximately unbiased test

To assess alternative relationships among *Lepidodinium chlorophorum*, MGD, and TGD in the 28S rRNA phylogeny (*SI Appendix*, Fig.S8), the trees with the highest likelihood were reconstructed from 28S rRNA alignment under four different topological constraints, namely (i) the monophyly of MGD and TGD, (ii) the monophyly of *L. chlorophorum* and MGD, (iii) the monophyly of *L. chlorophorum* and TGD, and (iv) the monophyly of the three species. The ML tree, in which the three species were paraphyletic, and the four alternative trees were subjected to an approximately unbiased test 60. For each test trees, site-wise log-likelihoods were calculated by RAxML with the GTR + Γ model. The resultant site-wise log-likelihood data were subsequently analyzed by Consel ver. 0.1 with default settings (73).

### Pigment analysis

For *L. chlorophorum*, we used strain NIES-1868 deposited in the National institute for Environmental studies (NIES) culture collection. MGD, TGD and *L. chlorophorum* cells were harvested by centrifugation. After removal of a transparent viscosity layer on the cell pellet, pigments were extracted with 100 µl of 100 % methanol, and the pigment extract was collected into a 1.5 ml centrifuge tube after centrifugation. We repeated the extraction until the extract was no longer colored. The extracted pigments were subjected to reverse-phase high-performance liquid chromatography (HPLC) with a Waters Symmetry C8 column (150 mm x 4.6 mm; particle size 3.5 µm; pore size 100 Å). The HPLC was performed as described in Zapata et al. (2000) (74) without any modifications. The eluted pigments were detected by the absorbance at 440 nm and identified by their elution patterns compared to those reported in Zapata et al. (2000) (74).

## Author Contributions

CS, KT, KI and MI established cultures and obtained morphological data. GT and TN carried out molecular analyses. CS, RK, HM performed HPLC analyses. GT, TN, RK, CS and YI drafted the manuscript. YI and GT designed this project. All authors read and approved the final manuscript.

## Acknowledge

This work was supported in part by grants from Japan Society for the Promotion of Sciences (23117006, 16H04826, 18KK0203 and 19H03280, YI; 25304029 and 15H04533, MI; 17H03723 and 26840123, GT; 14J05929, CH; 17K15164, TN; 19H03274, RK), Ministry of Education, Culture, Sports, Science and Technology of Japan Grant-in-Aid for Scientific Research on Innovative Areas 3308, a research grant from The Yanmar Environmental Sustainability Support Association for RY, and the “Tree of Life” research project of the University of Tsukuba.

## Supplementary Appendix

Fig. S1 HPLC chromatograms of pigments extracted from strains MGD and TGD as well as *Lepidodinium chlorophorum* strain NIES-1868. X1-X3 were explicit peaks of unknown pigments.

Fig. S2 Transmission electron microscopy image of an isolated plastid of strain TGD, showing the nucleomorph (Nm), plastid (Pl) and periplastidal compartment (PPC). Scale bar = 500 nm.

Fig. S3 Gene expression level of the green algal and alveolate transcripts. Natural logarithms of FPKM (fragments per kilo-base transcript length per million fragments mapped), which show putative abundances of transcripts, are indicated by color of markers in GC% scatter plots.

Fig. S4 Reverse transcription PCR using a dinoflagellate spliced-leader sequence primer and gene-specific primers. For TGD, we examined the 5′ termini of six ‘green algal transcripts’ encoding light-harvesting complex (LHC; lane 1), glutamyl-tRNA reductase (HemA; lane 2), eukaryotic initiation factor-4A (eIF4A; lane 3), plastocyanin (PetE; lane 4), ferredoxin (PetF; lane 5), and cytochrome *b6/f* complex iron-sulfur subunit (PetC; lane 6) in the upper image. For MGD, we examined the 5 ′ termini of six transcripts encoding ATP-dependent RNA helicase (lane 1), small subunit ribosomal protein S1 (RP-S1; lane 2), pyrophosphate-fructose 6-phosphate 1-phosphotransferase (PFK; lane 3), plastocyanin (PetE; lane 4), photosystem I subunit II (PsaD; lane 5), and cytochrome *b6/f* complex iron-sulfur subunit (PetC; lane 6) in the lower image.

Fig. S5 G + C contents (GC%) of green algal ‘house-keeping’ and ‘photosynthesis-related’ genes. In these scatter plots, the distributions of GC% of green algal transcripts are indicated as dots (see also Figs. 2a and 2b). In (a) and (b), red dots highlight the transcripts encoding house-keeping proteins in MGD and TGD, respectively. Definition of house-keeping proteins were adopted from previous studies on the nucleomorph genomes (references), including ribosomal proteins, transcription related factors, RNA polymerase subunits and spliceosomal subunits. The plots (c) and (d) are same as (a) and (b), but highlight transcripts encoding proteins involved in a different functional category. Blue dots represent the transcripts encoding photosynthesis-related proteins involved in photosystems I and II, cytochrome *b6*/*f*, light harvesting complexes and plastidal ATP synthesis in MGD (c) and TGD (d).

Fig. S6 The examples for SignalP predictions of green algal proteins with putative bipartite signal.

Fig. S7 Maximum-likelihood (ML) tree inferred from plastidal small subunit ribosomal RNA (16S rRNA) sequences with *Lepidodinium chlorophorum*. Only ML bootstrap support values greater than 80% are shown on the corresponding branches. The branches supported by Bayesian posterior probabilities greater than 0.95 are shown as thick lines.

Fig. S8 Maximum-likelihood (ML) tree inferred from eukaryotic large subunit ribosomal RNA (28S rRNA) sequences. Only ML bootstrap support values greater than 80% are shown on the corresponding branches. The branches supported by Bayesian posterior probabilities greater than 0.95 are shown.

