## Supplementary materials for "Dinoflagellates with relic endosymbiont nuclei as novel models for elucidating organellogenesis"

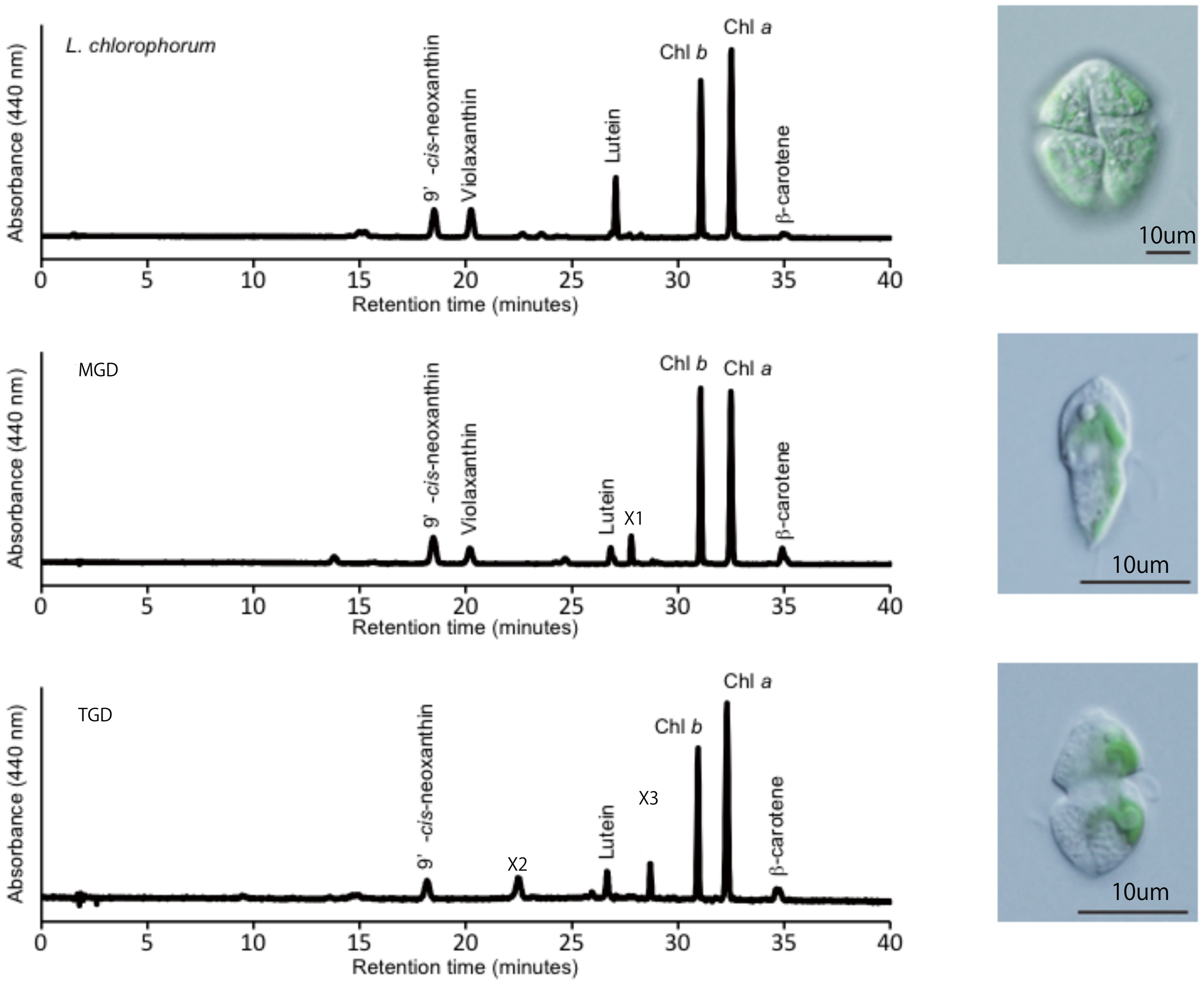

L. chlorophorum
MGD
TGD
Absorbance (440 nm)
Retention time (minutes)
9' -cis-neoxanthin
Violaxanthin
Lutein
X1
X2
X3
Chl b
Chl a
β-carotene
0
5
10
15
20
25
30
35
40
10um

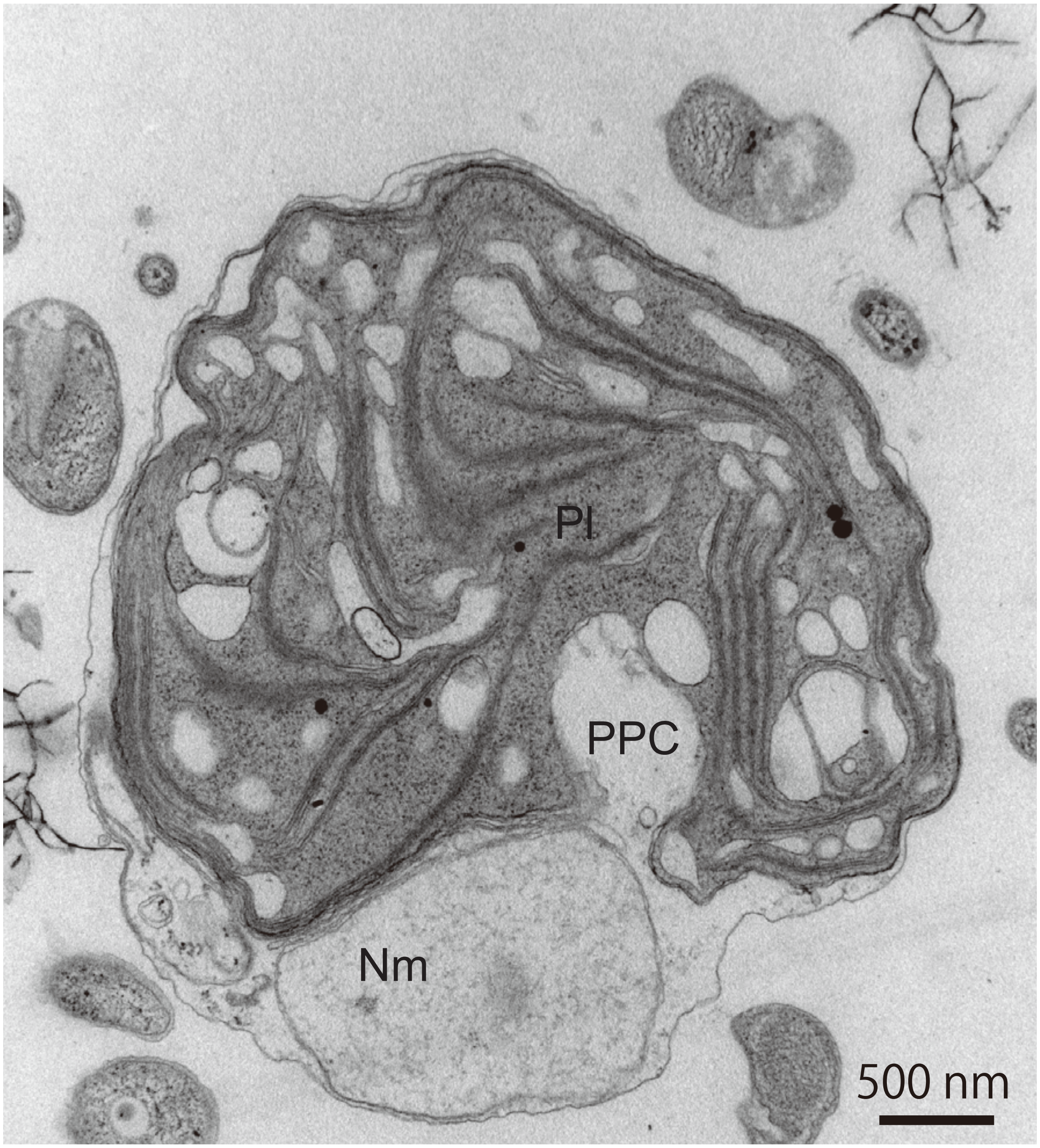
Pl
PPC
Nm
500 nm

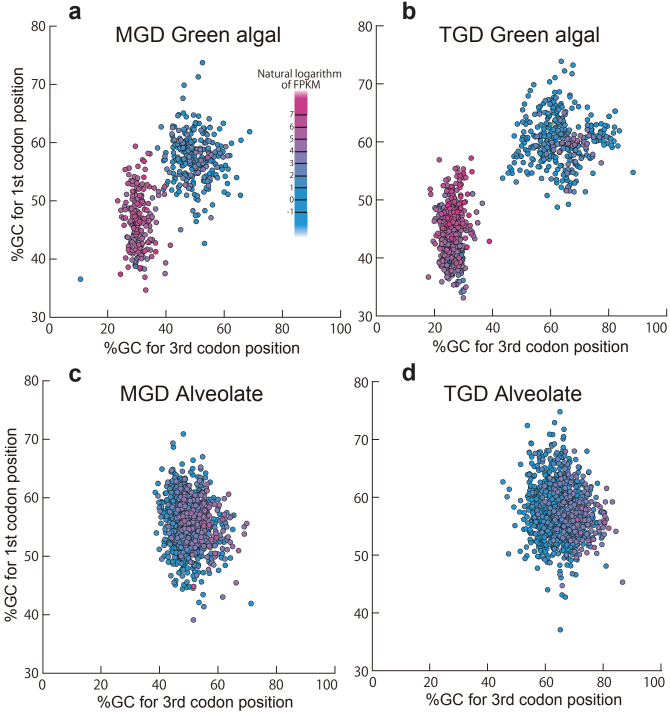

a
MGD Green algal
Natural logarithm of FPKM
7
6
5
4
3
2
1
0
-1
%GC for 1st codon position
%GC for 3rd codon position
b
TGD Green algal
c
MGD Alveolate
d
TGD Alveolate

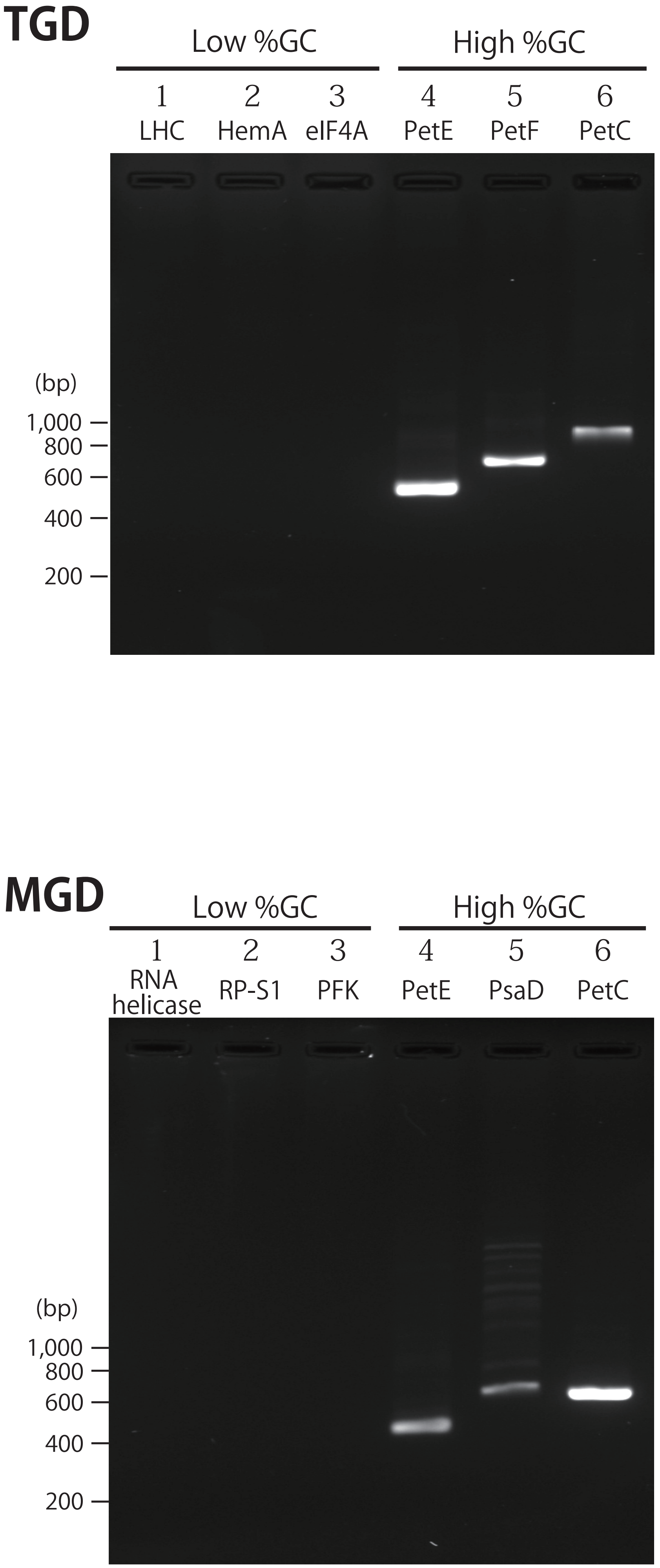
TGD
Low %GC
High %GC
1 LHC
2 HemA
3 eIF4A
4 PetE
5 PetF
6 PetC
(bp)
1,000
800
600
400
200
MGD
Low %GC
High %GC
1 RNA helicase
2 RP-S1
3 PFK
4 PetE
5 PsaD
6 PetC
(bp)
1,000
800
600
400
200

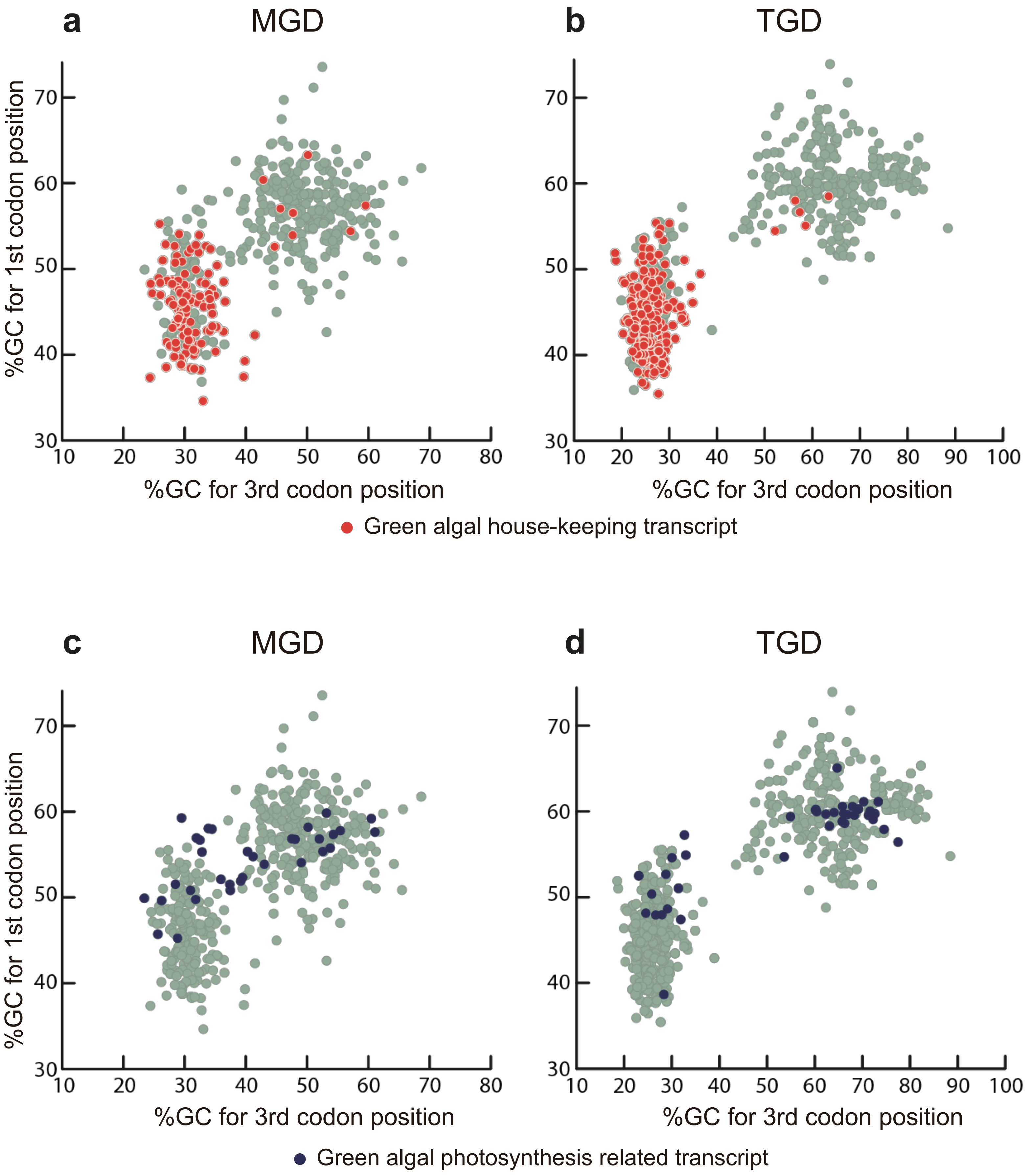
a
MGD
b
TGD
%GC for 1st codon position
%GC for 3rd codon position
Green algal house-keeping transcript
c
MGD
d
TGD
%GC for 1st codon position
%GC for 3rd codon position
Green algal photosynthesis related transcript

### Large subunit ribosomal protein L21 (plastid)

>MGD_42250_c0_seq1

MMRSIAALISFAYIASCAEQGDAQGSMNKLADKLADRVLN
VDPCTTDLENAVVAKGPGHLSSPTIQNRAPIMQPRGALSV
QSLARPVYCRSATEAAPWDAIIEVGGSQKIVETGRYYDTN
RLKGDLAPGTKVAFPRVLAVKKGAGEYTIGQPWVQGAKVE
AEVLENFKGKKVIVYKMRAKKHYRKKNGHRQLLTRFLVTN
VVKP

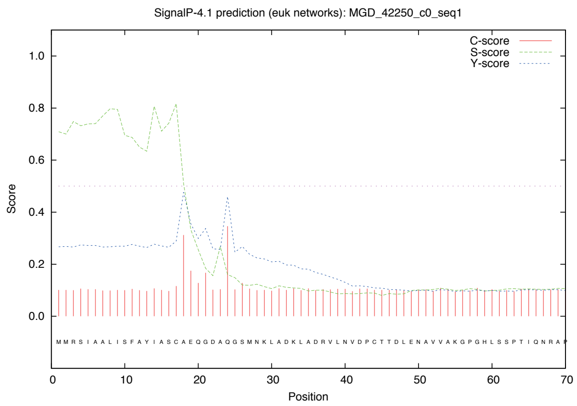


### Tocopherol O-methyltransferase

>MGD_65218_c0_seq1

MSSLLLMIIFVFTISWQAYTVNPQSLQGYHVGDTIDSLDT
LADRLANRLFDRVFKTKAMSPCALFSPSIKRWPHLLKKKI
PESGGNKWIVRSSRTREELNSGIAEFYDESSALWEGMWGE
HMHHGYYPGGKFRSDHQQAQIDMVDRVLDWAGVADVKNFL
DVGCGIGGSSRHIARRYGFPRHKISEKIARRYGLPLGVTG
QGITLSPNQAARANQLSSEQGLGEKLLFRVADALNMPFTD
GQFDLVWSLESGEHMPDKSQFLDELARVTAPGGK....

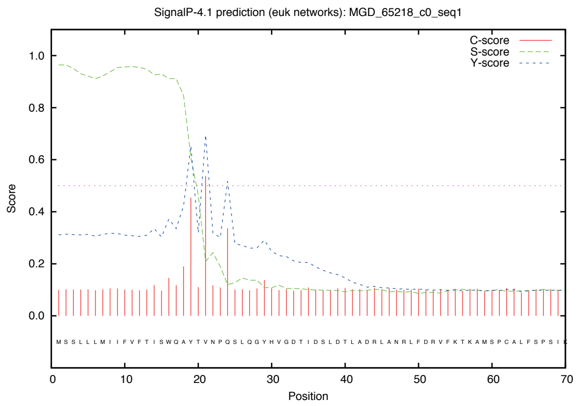


### Glucose-6-phosphate dehydrogenase

>MGD_149202_c0_seq1

MSTSRMCRASLNFLICCVAHGHVTNHSQAGWVGGLVDEFA
GQLVSKLFDRALDAPPLQRAELQRSMLAKPSHVAIPSHTH
VPLVPLKAHAHSFYSSGALRYQPQQGKAILAQHMRVGQTV
QKSLGNWARQLQSTPSRYADQLQAVDPDFLPSLYGADEDL
TLVIIGASGDLARKKVFPAIFALYAQGLLPKSTHIVGYAR
TDLSRDEFIERISEKLMCRIDWDAPDCSDDMDKFLSLTDY
VSGQYDSEADFAKLDAFITQKEVERKAKASNRLF....

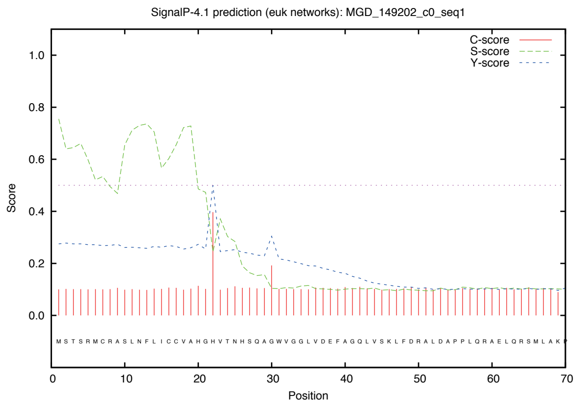


**RED:** Signal Peptide predicted by SignalP-4.1
**BLUE:** Putative mature protein region

### Photosystem II 13kDa protein (Psb28)

>TGD_36373_c0_seq1

MCCIRVAILLACIAQVSAKQTAADDAMDKLADRLVDKLAD
KLSDRLNQASSLHSADMDGTTLGKTSAIAAPQPRAVARAA
GVAPRMGMPFGMGAAGRVVQQRMPVLANAGASLQFIKGTD
EPDVPEVKLSKSRSSSMGQATFIFENPSVFDLEGPGKDDI
TGLYMVDDEGEMRTVDVQARFVNGKPAGIIAKYTMQNEAQ
WDRFMRFMERYAEANDLGFNKAK

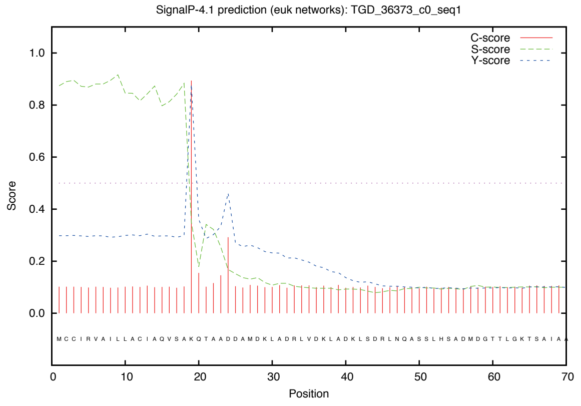


### 4-alpha-glucanotransferase

>TGD_140644_c0_seq2

MLKAAYILLASVSCVDANELVVHDAAFAPAFIDTVVDRLV
DKLAKRTRSSDLGSTTLGKAVHVATRCRPGSQPCALSTSH
SMPAWPRTYHVARPMTVPRQTADFEKIFAGVSTKRSLPVV
QAQQAEVAKEAFKFDLKRRAGVLLPVSSLDGQGPIGNLDD
AERFVDWLAEAGMALWQILPLVPTDSAGSPYSSWSTLSGN
PDLVGLGGLVAAGLLDKEKTKLPLLTTVNYTVVAAQKRTL
VLEAAQALLDRPDHPLRPALDKFVANAKWATDAA....

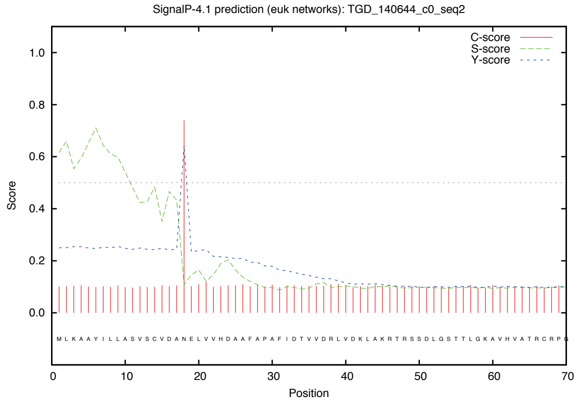


### Nucleoside-diphosphate kinase

>TGD_174950_c0_seq1

MMSKVAAAVLFVVVAQTFAQQTAVSQYDLANKVADKLAAK
LLDRMEDANLDDTTLGALLQSSNMQLAGLSPSTQLAFSAG
PLRTPLAPSRIALNAWESDTCARRAVSAAAVAGVPRMCAA
KEVRALGLRKSQPTHVVQASASAERSYVMIKPDGVQRGLV
GEIISRFERKGFYLKGLKMFQTPEDLAKEHYKDLSEKPFF
GDLVEYICSGPVVCMVWEGKGVIKSARKLIGATNPLEAEP
GTIRGDFAVETGRNVIHGSDSIENGEREIGIWF....

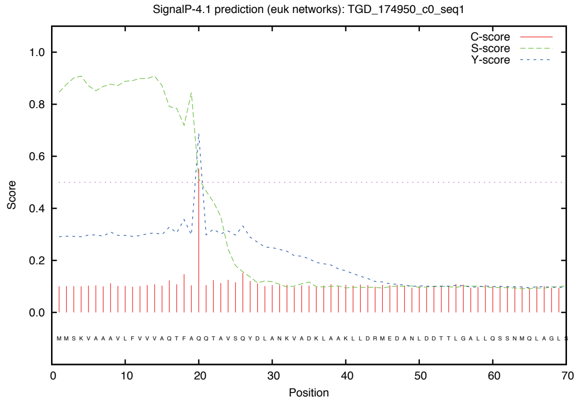


**RED:** Signal Peptide predicted by SignalP-4.1
**BLUE:** Putative mature protein region

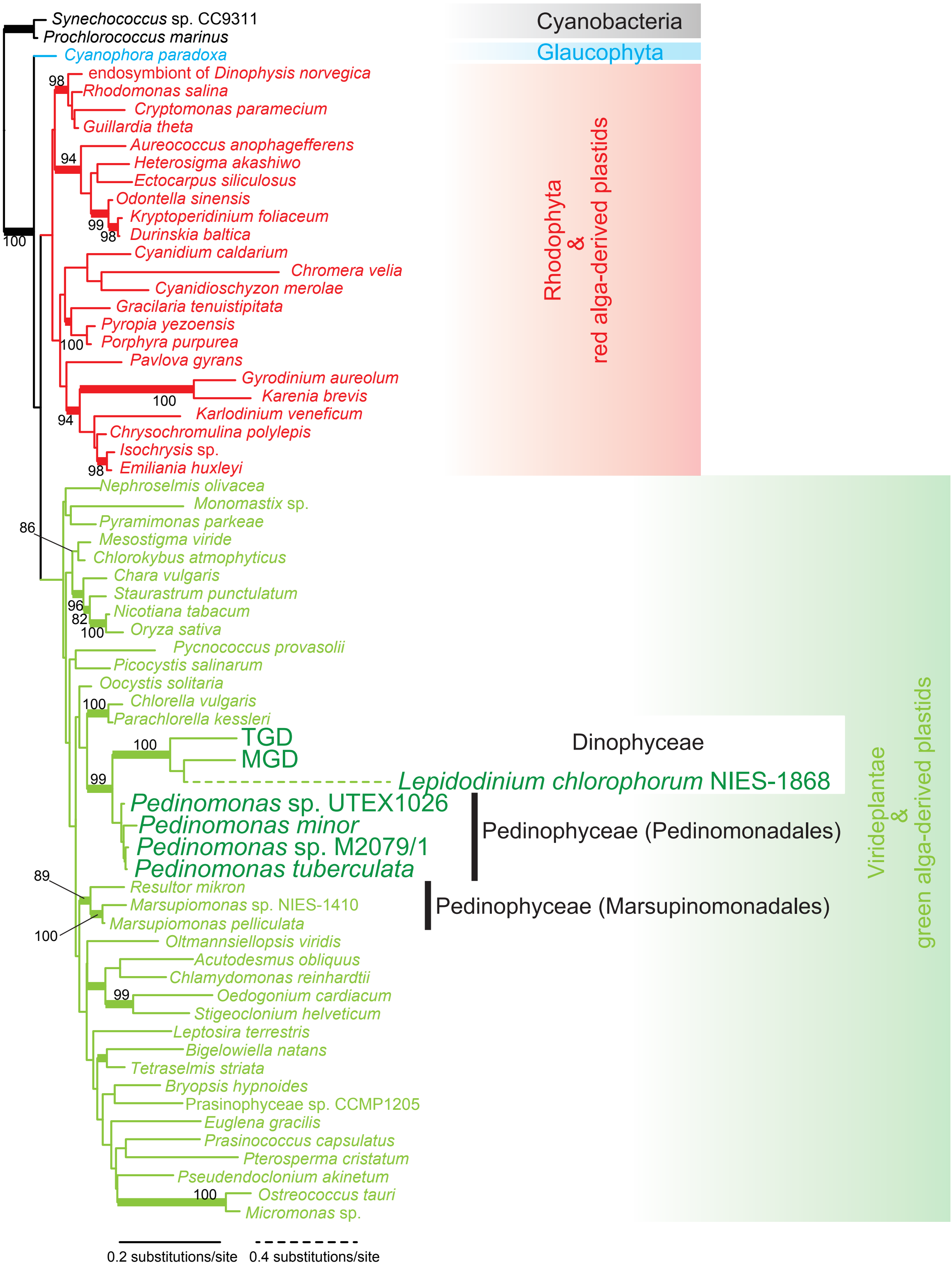

Synechococcus sp. CC9311
Prochlorococcus marinus
Cyanophora paradoxa
endosymbiont of Dinophysis norvegica
Rhodomonas salina
Cryptomonas paramecium
Guillardia theta
Aureococcus anophagefferens
Heterosigma akashiwo
Ectocarpus siliculosus
Odontella sinensis
Kryptoperidinium foliaceum
Durinskia baltica
Cyanidium caldarium
Chromera velia
Cyanidioschyzon merolae
Gracilaria tenuistipitata
Pyropia yezoensis
Porphyra purpurea
Pavlova gyrans
Gyrodinium aureolum
Karenia brevis
Karlodinium veneficum
Chrysochromulina polylepis
Isochrysis sp.
Emiliania huxleyi
Nephroselmis olivacea
Monomastix sp.
Pyramimonas parkeae
Mesostigma viride
Chlorokybus atmophyticus
Chara vulgaris
Staurastrum punctulatum
Nicotiana tabacum
Oryza sativa
Pycnococcus provasolii
Picocystis salinarum
Oocystis solitaria
Chlorella vulgaris
Parachlorella kessleri
TGD
MGD
Lepidodinium chlorophorum NIES-1868
Pedinomonas sp. UTEX1026
Pedinomonas minor
Pedinomonas sp. M2079/1
Pedinomonas tuberculata
Resultor mikron
Marsupiomonas sp. NIES-1410
Marsupiomonas pelliculata
Oltmannsiellopsis viridis
Acutodesmus obliquus
Chlamydomonas reinhardtii
Oedogonium cardiacum
Stigeoclonium helveticum
Leptosira terrestris
Bigelowiella natans
Tetraselmis striata
Bryopsis hypnoides
Prasinophyceae sp. CCMP1205
Euglena gracilis
Prasinococcus capsulatus
Pterosperma cristatum
Pseudendoclonium akinetum
Ostreococcus tauri
Micromonas sp.
Cyanobacteria
Glaucophyta
Rhodophyta & red alga-derived plastids
Dinophyceae
Pedinophyceae (Pedinomonadales)
Pedinophyceae (Marsupinomonadales)
Viridiplantae & green alga-derived plastids
0.2 substitutions/site
0.4 substitutions/site

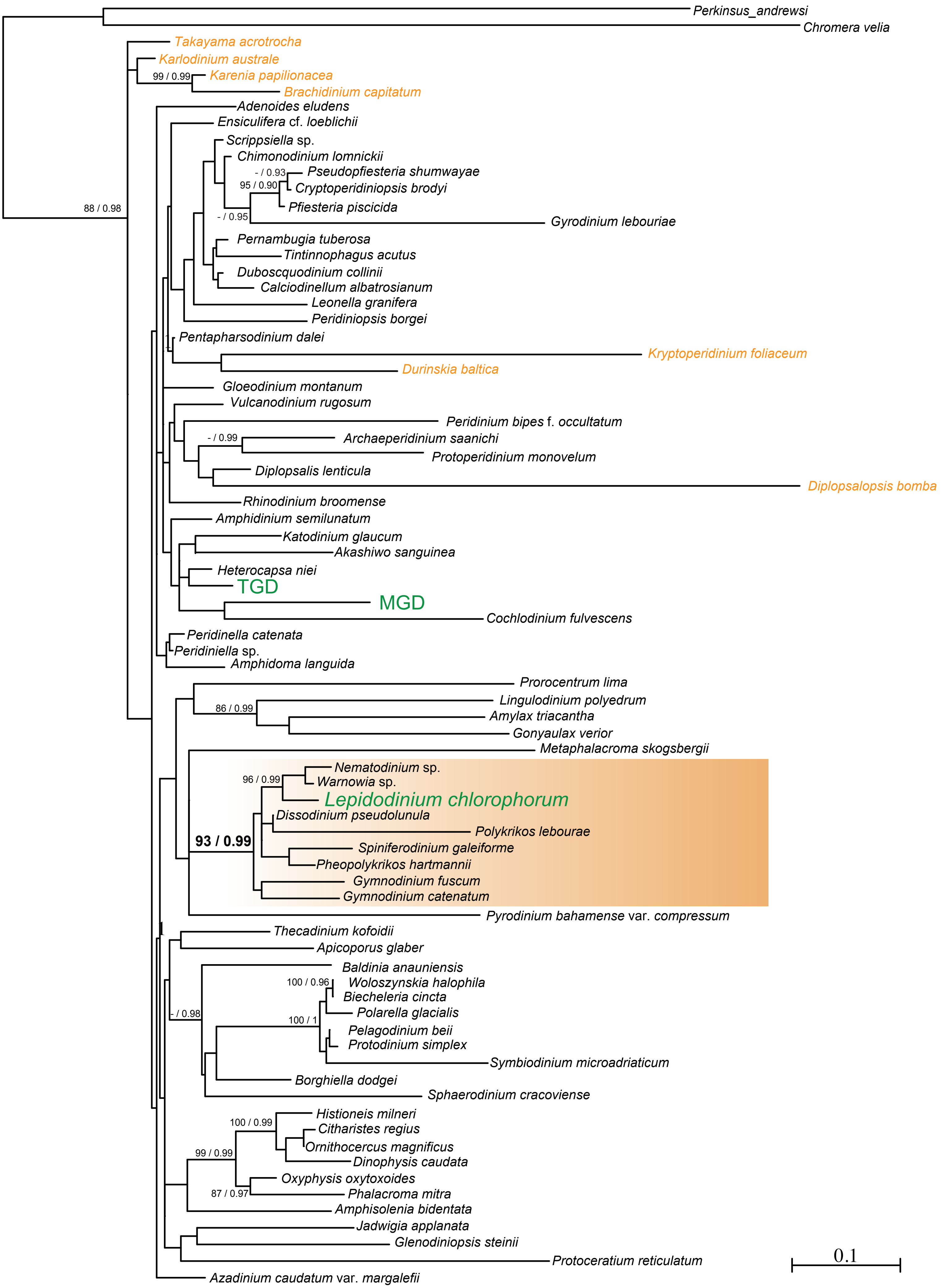
Perkinsus_andrewsi
Chromera velia
Takayama acrotrocha
Karlodinium australe
99 / 0.99
Karenia papilionacea
Brachidinium capitatum
88 / 0.98
Adenoides eludens
Ensiculifera cf. loeblichii
Scrippsiella sp.
Chimonodinium lomnickii
- / 0.93
Pseudopfiesteria shumwayae
95 / 0.90
Cryptoperidiniopsis brodyi
Pfiesteria piscicida
- / 0.95
Gyrodinium lebouriae
Pernambugia tuberosa
Tintinnophagus acutus
Duboscquodinium collinii
Calciodinellum albatrosianum
Leonella granifera
Peridiniopsis borgei
Pentapharsodinium dalei
Kryptoperidinium foliaceum
Durinskia baltica
Gloeodinium montanum
Vulcanodinium rugosum
Peridinium bipes f. occultatum
- / 0.99
Archaeperidinium saanichi
Protoperidinium monovelum
Diplopsalis lenticula
Diplopsalopsis bomba
Rhinodinium broomense
Amphidinium semilunatum
Katodinium glaucum
Akashiwo sanguinea
Heterocapsa niei
TGD
MGD
Cochlodinium fulvescens
Peridinella catenata
Peridiniella sp.
Amphidoma languida
Prorocentrum lima
86 / 0.99
Lingulodinium polyedrum
Amylax triacantha
Gonyaulax verior
Metaphalacroma skogsbergii
Nematodinium sp.
96 / 0.99
Warnowia sp.
Lepidodinium chlorophorum
Dissodinium pseudolunula
93 / 0.99
Polykrikos lebourae
Spiniferodinium galeiforme
Pheopolykrikos hartmannii
Gymnodinium fuscum
Gymnodinium catenatum
Pyrodinium bahamense var. compressum
Thecadinium kofoidii
Apicoporus glaber
Baldinia anauniensis
100 / 0.96
Woloszynskia halophila
Biecheleria cincta
- / 0.98
Polarella glacialis
100 / 1
Pelagodinium beii
Protodinium simplex
Symbiodinium microadriaticum
Borghiella dodgei
Sphaerodinium cracoviense
Histioneis milneri
100 / 0.99
Citharistes regius
99 / 0.99
Ornithocercus magnificus
Dinophysis caudata
Oxyphysis oxytoxoides
87 / 0.97
Phalacroma mitra
Amphisolenia bidentata
Jadwigia applanata
Glenodiniopsis steinii
Protoceratium reticulatum
0.1
Azadinium caudatum var. margalefii
